# Targeting of mitochondrial and cytosolic substrates of tRNA isopentenyltransferases: selection of differential tRNA-i6A37 identity subsets

**DOI:** 10.1101/714972

**Authors:** Abdul Khalique, Sandy Mattijssen, Alexander F. Haddad, Richard J. Maraia

## Abstract

tRNA isopentenyltransferases (IPTases), which add an isopentenyl group to *N*^*6*^ of adenosine-37 (i6A37) of certain tRNAs, are among a minority of modification enzymes that act on both cytosolic and mitochondrial substrates. The *Caenorhabditis elegans* mitochondrial IPTase impacts life expectancy, and pathogenic mutations to human IPTase (TRIT1) that decrease i6A37 levels cause mitochondrial insufficiency and neurodevelopmental disease. Understanding of IPTase broad function should consider the differential identities of the tRNAs selected for i6A37 formation and their cognate codons, which vary among species in both their nuclear- and mitochondria-encoded tRNAs. Substrate selection is principally by recognition of the A36-A37-A38 sequence but can be negatively impacted by certain anticodons, and by ill-defined properties of the IPTase. Thus, tRNAs-i6A37 comprise a modification code system whose principles are incompletely understood. While *Saccharomyces cerevisiae* uses alternative translation initiation to target IPTase to mitochondria, our analyses indicate that TRIT1 uses a single initiation site to produce a mitochondrial targeting sequence (MTS) that we demonstrate by point mutagenesis using GFP imaging in human cells. We also examined cytosolic and mitochondrial tRNA modification by TRIT1 in *Schizosaccharomyces pombe* using tRNA-mediated suppression and i6A37-sensitive northern blotting. The TRIT1 MTS mutations indeed decrease mitochondrial-tRNA modification in *S. pombe*. We also show TRIT1 modification deficiency specific for tRNA^**Trp**^CCA despite A36-A37-A38, consistent with the negative effect of the CCA anticodon as was described for Mod5 IPTase. This TRIT1 deficiency can be countered by over-expression. We propose a model of tRNA-i6A37 identity selection in eukaryotes that includes sensitivity to substrates with YYA anticodons.

**AUTHOR SUMMARY:** tRNA isopentenyltransferases (IPTases) are tRNA modification enzymes that are conserved in bacteria and eukaryotes. They add an isopentenyl group to the Adenosine base at position 37, adjacent to the anticodon of specific subsets of tRNAs that decode codons that begin with Uridine. This modification stabilizes the otherwise weak adjacent codon-anticodon basepair and increases the efficiency of decoding of the corresponding codons of the genetic code. IPTases belong to a group of enzymes that modify both cytoplasmic and mitochondrial tRNAs of eukaryotic cells. Interestingly, during evolution there were changes in the way that IPTases are targeted to mitochondria as well as changes in the relative numbers and identities of IPTase tRNA substrates in the cytoplasm vs. mitochondria, the latter consistent with phenotypic consequences of IPTase deficiencies in fission and budding yeasts, and mammals. Pathogenic mutations to human IPTase (TRIT1) cause mitochondrial insufficiency and neurodevelopmental disease, principally due to decreased modification of the mt-tRNA substrates. In this study, we identify the way human TRIT1 is targeted to mitochondria. We also show that TRIT1 exhibits a tRNA anticodon identity-specific substrate sensitivity. The work leads to new understanding of the IPTases and the variable codon identities of their tRNA substrates found throughout nature.

## INTRODUCTION

45 different eukaryotic cytoplasmic (cy-) tRNAs contain unique sets of modifications in two groups [1-3]; those in the body generally contribute to folding and stability, and those in the anticodon loop (ACL) contribute to mRNA decoding by ribosomes. One ACL modification is isopentenyl addition to *N*^*6*^ of A37 (i6A37), immediately 3’ to anticodons that decode UNN codons. The i6A37 and/or its derivative (below) has been shown to increase anticodon-codon binding efficiency, promote mRNA decoding, and decrease frameshifting [reviewed in 4]. Modifications in the anticodon stem loop (ASL) are more concentrated and diverse relative to the tRNA body, and are involved in modification circuits [5], some of which are disrupted by mutations that cause diseases [6, 7]. In one ASL circuit, i6A37 is prerequisite for m3C32 formation by Trm140 and related enzymes [8, 9].

The tRNA isopentenyltransferases (IPTases) that form i6A37 have been highly conserved and characterized from human (TRIT1), fission yeast *S. pombe* (Tit1/Sin1), budding yeast *S. cerevisiae* (Mod5), roundworm *C. elegans* (GRO-1), bacteria *Escherichia coli* (MiaA), and others [4]. IPTases are members of a minority group of modification enzymes that act on both cytoplasmic (cy)-tRNAs and mitochondrial (mt)-tRNAs [10, 11]. Bacterial i6A37 are mostly hypermodified to 2-methylthio-*N*^6^-isopentenyl-A37 (ms2i6A37), whereas eukaryotic cy-tRNAs i6A37 are not. While the mt-tRNAs of *S. cerevisiae* and *S. pombe* are not hypermodified, the mammalian mt-tRNAs i6A37 are [12-14].

Identities of tRNAs with i6A37 vary remarkably in nature. The number of cy-tRNAs-i6A37 is highest in bacteria, intermediate in yeast and lowest in human (Table 1) [15]. The wide variable identities of tRNAs that harbor i6A37 among species is in stark contrast to the threonylcarbamoyl-A37 (t6A37) found on all cy-tRNAs that decode ANN codons in all species (reviewed in [16], also see [17]). Also notably, tRNA-i6A37 identities vary highly between cytosol and mitochondria among and within species (Table 1). Much of the documented variability in cy-tRNA-i6A37 identity is reflected by loss of the IPTase recognition element, A36-A37-A38; as evolutionary complexity increased, position 37 was occupied more often by G than A in cy-tRNAs that decode UNN codons [16]. However, this doesn’t account for all of the variability; absence of i6A37 from *S. cerevisiae* cy-tRNA^**Trp**^CCA, which bears A36-A37-A38, reflects substrate-specific deficiency of Mod5 due to poor recognition of or negative influence of the CCA anticodon [17]. *S. pombe* Tit1, which differs from Mod5 in the number and composition of amino acids in its anticodon recognition loop motif, can readily modify the identical tRNA^**Trp**^CCA [17]. Thus, different lines of evidence indicate that two mechanisms have contributed to the species-specific profiles of tRNA-i6A37 identities in these model organisms and possibly others. This led to the idea that the i6A37 modification system represents a plastic component of the genetic code [18] and also suggests its adaptability for synthetic biology [19]. Insights into such potential will require understanding of the elements of differential substrate targeting by the IPTases. Yet, the principles of tRNA identity-recognition for i6A37 selection are incompletely understood.

**Table 1.**

| tRNA anticodon<br>(as set by encoding<br>DNA) | *E. coli*<br>*MiaA* | *S. pombe*<br>*tit1*<sup>+</sup> | *S. cerevisiae*<br>*MOD5* | human or bovine<br>*TRIT1* |
| --- | --- | --- | --- | --- |
| cy-Ser GGA | A<sub>37</sub> | — | — | — |
| cy-Ser AGA | — | i<sup>6</sup>A<sub>37</sub> | i<sup>6</sup>A<sub>37</sub> | i<sup>6</sup>A<sub>37</sub> |
| cy-Ser CGA | ms<sup>2</sup>i<sup>6</sup>A<sub>37</sub> | i<sup>6</sup>A<sub>37</sub> | i<sup>6</sup>A<sub>37</sub> | i<sup>6</sup>A<sub>37</sub> |
| cy-Ser UGA | ms<sup>2</sup>i<sup>6</sup>A<sub>37</sub> | i<sup>6</sup>A<sub>37</sub> | i<sup>6</sup>A<sub>37</sub> | i<sup>6</sup>A<sub>37</sub> |
| cy-Ser<sup>Sec</sup> UCA | i<sup>6</sup>A<sub>37</sub> | — | — | i<sup>6</sup>A<sub>37</sub> |
| cy-Tyr GUA | ms<sup>2</sup>i<sup>6</sup>A<sub>37</sub> | i<sup>6</sup>A<sub>37</sub> | i<sup>6</sup>A<sub>37</sub> | m<sup>1</sup>G<sub>37</sub> |
| cy-Cys GCA | ms<sup>2</sup>i<sup>6</sup>A<sub>37</sub> | G | i<sup>6</sup>A<sub>37</sub> | m<sup>1</sup>G<sub>37</sub> |
| cy-Trp CCA | ms<sup>2</sup>i<sup>6</sup>A<sub>37</sub> | i<sup>6</sup>A<sub>37</sub> | A<sub>37</sub> | m<sup>1</sup>G<sub>37</sub> |
| cy-Phe GAA | ms<sup>2</sup>i<sup>6</sup>A<sub>37</sub> | yW<sub>37</sub> | yW<sub>37</sub> | yW<sub>37</sub> |
| cy-Leu CAA | ms<sup>2</sup>i<sup>6</sup>A<sub>37</sub> | G | m<sup>1</sup>G<sub>37</sub> | m<sup>1</sup>G<sub>37</sub> |
| cy-Leu UAA | ms<sup>2</sup>i<sup>6</sup>A<sub>37</sub> | G | m<sup>1</sup>G<sub>37</sub> | m<sup>1</sup>G<sub>37</sub> |
| **# of i<sup>6</sup>A cy-cognate codons** | **12** | **7** | **8** | **5** |
| mt-Trp CCA<sup>#</sup> |  | i<sup>6</sup>A<sub>37</sub> | — | — |
| mt-Trp UCA<sup>#</sup> |  | — | i<sup>6</sup>A<sub>37</sub> | ms<sup>2</sup>i<sup>6</sup>A<sub>37</sub> |
| mt-Tyr GUA |  | G | i<sup>6</sup>A<sub>37</sub> | ms<sup>2</sup>i<sup>6</sup>A<sub>37</sub> |
| mt-Ser UGA |  | G | m<sup>1</sup>G<sub>37</sub> | ms<sup>2</sup>i<sup>6</sup>A<sub>37</sub> |
| mt-Phe GAA |  | G | m<sup>1</sup>G<sub>37</sub> | ms<sup>2</sup>i<sup>6</sup>A<sub>37</sub> |
| mt-Cys GCA |  | i<sup>6</sup>A<sub>37</sub> | i<sup>6</sup>A<sub>37</sub> | i<sup>6</sup>A<sub>37</sub> |
| mt-Leu UAA |  | A<sup>*</sup> | m<sup>1</sup>G<sub>37</sub> | A<sup>*</sup> |
| **# of i<sup>6</sup>A mt-cognate codons** |  | **4** | **6** | **13** |

<sup>#</sup>Position 34 of *S. pombe* mt-tRNA<sup>Trp</sup> is encoded as C [91] whereas in *S. cerevisiae* [92, 93], bovine and human [94], it is encoded to be U, and was determined to be 5-taurinomethyluridine (tm<sup>5</sup>U) in bovine [12], and cmnm<sup>5</sup>U in *S. cerevisiae* [85] (see tRNAmoViz server [84]). All entries with subscript <sub>37</sub> have been verified
as modified or unmodified; entries without <sub>37</sub> show encoded nucleotide but have not been validated [12, 17, 42, 64]. *S. cerevisiae* mt-tRNA<sup>Trp</sup> i6A37 was reported in [95]. \* A is not in an A36-A37-A38 context. yW: ybutosine; dash “-” indicates there is no gene for a tRNA with this anticodon [81]. *Sup-3e* [25] (encodes cy-tRNA<sup>Ser</sup>UCA, also referred to as pSer tRNA [32, 96]) contains mcm<sup>5</sup>U [86] as does mammalian selenocysteine-inserting cy-tRNA<sup>Ser[Sec]</sup>UCA [87]. See *E. coli* tRNA<sup>Ser[Sec]</sup>UCA [97]. The mt-tRNAs<sup>Trp</sup> recognize 2 sense codons in yeast and animal mitochondria [98] [12].

**Table 2.**
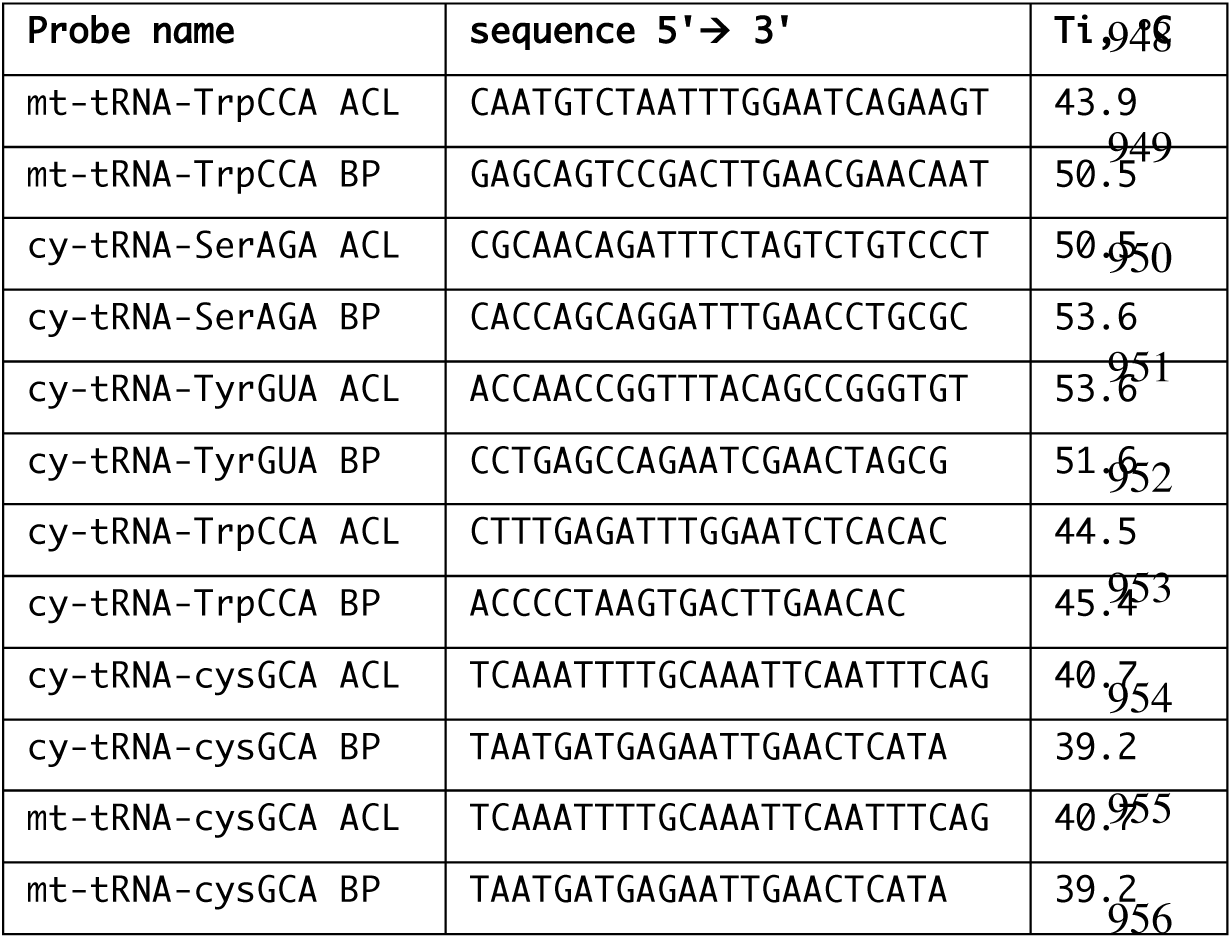
Oligo-DNAs used for PHA6 assay; ACL (anticodon loop), BP (body probe).

**Table 3.**
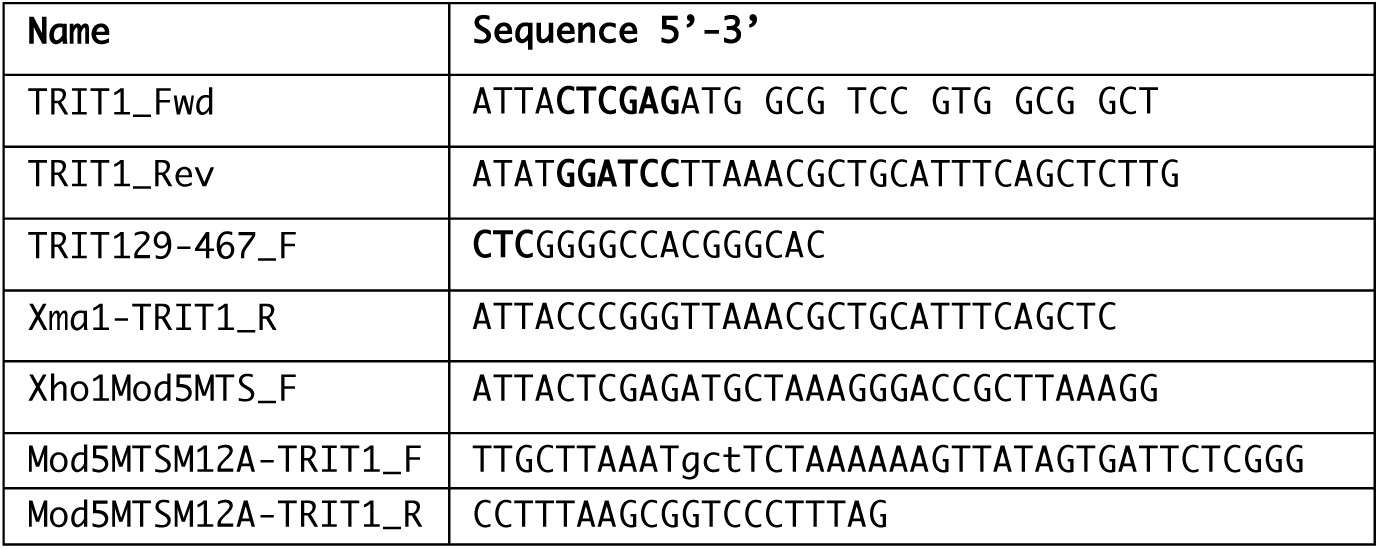
PCR Primers used in this study

Work on *S. pombe* and *S. cerevisiae* IPTases used suppressor-tRNAs^**Ser**^UCA and ^**Tyr**^UAA and a red -white assay based on decoding premature stop codons in mRNAs for adenine synthesis [20-22]. tRNA-mediated suppression (TMS) can be used to study many facets of tRNA biogenesis [23-32].

Human TRIT1 was identified as a tumor suppressor [33], and an allele that encodes TRIT1-F202L was later found to be associated with modulation of lung cancer survival [34] [also see 35, 36]. IPTases use dimethylallyl pyrophosphate (DMAPP) for i6A37 formation, a substance also used for sterol synthesis [37], relevant because Mod5 prions inactive for modification, support increased sterol metabolism and anti-fungal drug resistance [38, 39]. Complementation of *S. cerevisiae* cy-tRNA^**Tyr**^UAA-TMS as a test of IPTases [33, 40, 41], showed TRIT1-F202L as active [34]. Using *S. pombe*, substitution of invariant Thr-12 in the P-loop of Tit1 (Fig 1, Tit1-T12A) inactivated it for cy-tRNA^**Ser**^-TMS [4, 17]. Tit1 T12A is homologous to the T19A substitution of *E. coli* IPTase which resulted in the largest decrease in *k*_*cat*_ (600-fold) of all mutations tested [17, 42, 43]. The corresponding T23 in Mod5 is indeed positioned for catalysis [44]. Thus, an outstanding puzzle has been that a TRIT1(57-467) variant missing the P-loop entirely as well as D55 (of DSMQ, Fig 1)), which in Mod5 acts as a general base for catalysis [44], complemented cy-tRNA^**Tyr**^-TMS [33]. Another enigma is a predicted proteolytic cleavage site upon mitochondrial targeting which would also remove the N-terminal region from TRIT1. Both of these are addressed by new data below.

**Figure 1.**
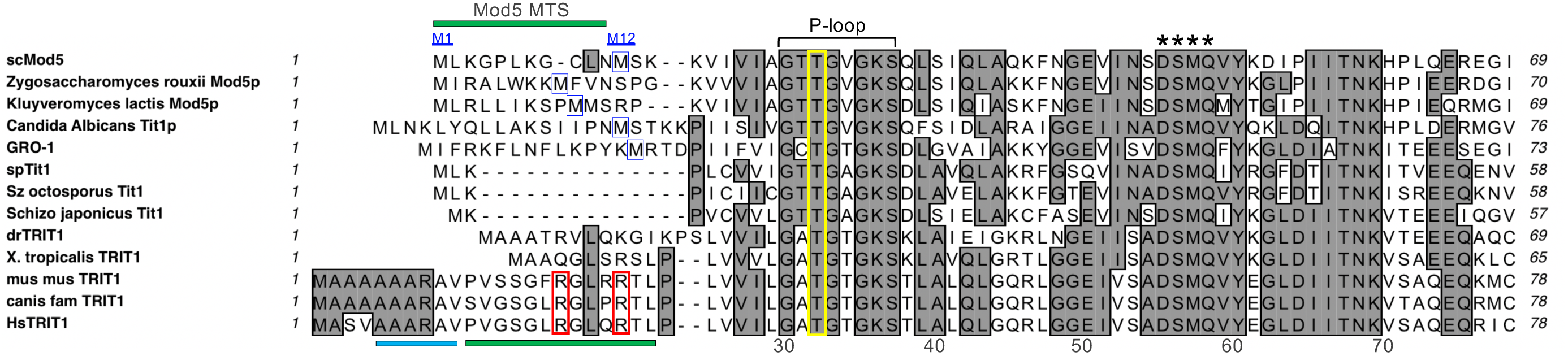
N-termini sequence alignment of representatives of phylogenetic groups of IPTases. PTase from *C. elegans*; spTit1: *S. pombe*; Sz = *S. octosporus*; dr: *D. rerio; mus mus: M. musculus; canis fam: C. familiaris Hs: H. sapiens*. The region necessary as a mitochondrial targeting sequence (MTS) of Mod5 is indicated [45]. The second methionine that may be used as alternative translation start sites for cytoplasmic localization are boxed in blue. The invariant Threonine within the conserved P-loop that functions in catalysis is in the yellow rectangle. Asterisks indicate the conserved DSMQ sequence that forms a network of contacts that mediate A37 recognition; the D of which in Mod5 acts as a general base for catalysis [44] and the M of which is M57 of TRIT1(57-467). As predicted by MitoFates [56], the TOM20-binding site of human TRIT1 is underlined with a blue bar and the amphipathic helices of TRIT1 and Mod5 are indicated with green bars. The two key basic residues in the amphipathic helices of the conserved mammalian MTSs that were mutated in human TRIT1 are in red rectangles.

Proper subcellular trafficking is critical for IPTase function; in yeast, the single nuclear gene-encoded Mod5 localizes to nuclei, cytoplasm and mitochondria [45-47]. Nuclear Mod5 is involved in tRNA gene-mediated silencing [48, 49]; its nucleolar colocalization [47] is consistent with this and with transcriptionally active tRNA genes and early tRNA processing [50, 51]. For some tRNAs, A36-A37-A38 is interrupted by an intron, and in yeast where tRNA splicing occurs in the cytoplasm, tRNA nuclear export is prerequisite to i6A37 formation [52]. However, tRNA^**Ser**^AGA has no intron and may be modified by nuclear Mod5. The Mod5 nuclear localization signal (NLS) is near its C-terminus [47], following a Zn-finger motif specific to eukaryotic IPTases [4] that binds the tRNA backbone [44], which has been shown to be required for Tit1 activity [17].

Modification of mt-tRNA is a very important IPTase function [12]. Mt-IPTase deficiency is linked to *C. elegans* longevity [53]. Mutations in TRIT1 cause neurodevelopmental disorder [41, 54]. In this case, most if not all pathophysiology is attributable to hypomodification of mt-tRNA and impairment of mitochondrial translation and oxidative phosphorylation, even though the cy-tRNAs are also hypomodified [41]. Human mt-tRNAs-i6A37 decode 13 codons for 5 amino acids in mitochondria whereas cy-tRNAs-i6A37 decode 5 codons for 1 amino acid in the cytosol (Table 1).

*S. cerevisiae* and *C. elegans* use alternative translation initiation to differentially target their IPTases to mitochondria and cytoplasm [45, 46, 53]. Other IPTase mRNAs do not use alternate starts and their distribution mechanisms have not been scrutinized. We show that TRIT1 contains an N-terminal MTS that functions in human and *S. pombe* cells. We employed TMS and i6A37-sensitive northern blotting to show that TRIT1 modifies cy-and mt-tRNAs in *S. pombe*. Our data also reveal severe deficiency of TRIT1 for cy-tRNA^**Trp**^CCA specifically, despite the A36-A37-A38 sequence, consistent with negative effects of the CCA anticodon as was described for Mod5 IPTase. We present a model of eukaryotic i6A37 modification based on enzyme concentration and substrate preference that extends to conditional YYN codon recognition, includes differential subcellular localization, and suggests a role for prerequisite influence by U34 and C34 modification.

## RESULTS

### TRIT1 bears a mitochondrial targeting sequence (MTS) but may not undergo cleavage

As noted previously [41], mitochondrial targeting of human TRIT1 was predicted with high confidence by the online tool, MitoProt II (http://ihg.gsf.de/ihg/mitoprot.html) [55], which also predicted a proteolytic cleavage site after amino acid 47. However, such cleavage would remove the highly conserved P-loop likely important for catalytic activity (Fig 1). We also used the prediction algorithms MitoFates [56] and iPSORT [57] to analyze the human TRIT1 sequence. iPSORT predicted a MTS within the first 30 amino acids but doesn’t predict cleavage. MitoFates was designed to be an improved method for predicting cleavable N-terminal MTSs, referred to as presequences, and their cleavage sites [56]. MitoFates predicted a MTS comprised of a TOM20 (translocase of the outer mitochondrial membrane-20) receptor binding site at amino acids 5-10 (AAARAV, Fig 1) followed by an amphipathic helix at 11-23 (PVGSGLRGLQRTL), but with a low probability of a cleavable presequence of 0.106, well below MitoFates default cutoff of 0.5. Low probability of a cleavable presequence is consistent with biochemical subfractionation that revealed no evidence of size difference between lysate and matrix-localized TRIT1 in a gel that revealed size difference for matrix-localized GDH [41]. We note that the cNLS mapper [58] predicted overlapping mono- and bi-partite importin-dependent nuclear localization signals at 420-453 of TRIT1.

We created GFP fusion constructs to test for mitochondrial localization by confocal microscopy in human embryonic kidney (HEK)-293 cells (Fig 2A, B). To test for MTS activity of the TRIT1 N-terminal region we fused only its first 51 amino acids to GFP, as in TRIT1(1-51)-GFP, and examined it for mitochondrial targeting. Two point mutations, R17E and R21E, in the basic amphipathic helix component of the predicted MTS (Fig 1) were made in two contexts, the N-terminal fusion 1-51 construct, TRIT1(1-51:R17E/R21E)-GFP, and full length TRIT1(R17E/R21E)-GFP. These and other constructs were transfected into HEK293 cells. After 48 hr the cells were fixed and exposed to MitoTracker to stain mitochondria red and with Hoechst to stain nuclei blue (Fig 2B). Cell lysates were examined for protein expression by western blot using anti-GFP antibody (Fig 2C). GFP alone localized diffusely in cytosol and nuclei consistent with diffusion for its mass of ∼25 kDa (Fig 2B, column 1). Full length TRIT1-GFP was found localized to nuclei and cytosol with some evidence of mitochondrial localization reflected by overlap of GFP and MitoTracker as a stippling yellow-orange pattern in the merge (Fig 2B, column 2, lower panel). This pattern was not observed with full length TRIT1(R17E/R21E)-GFP containing the MTS mutations, which accumulated in nuclei and was not found to localize to mitochondria (Fig 2B, column 3). We note from these results that most cytoplasmic TRIT1 would appear to be due to mitochondrial localization (Discussion). The fragment lacking the first 51 amino acids, TRIT1(52-467)-GFP, which did not accumulate to the same levels as the other proteins (Fig 2C). This fusion protein was found expressed in fewer transfected cells than the other constructs and although we do not know the basis of this, some cells appeared to contain aggregates of GFP (data not shown). In cells in which aggregates were not observed, TRIT1(52-467)-GFP was accumulated in nuclei (Fig 2B, column 6).

**Figure 2.**
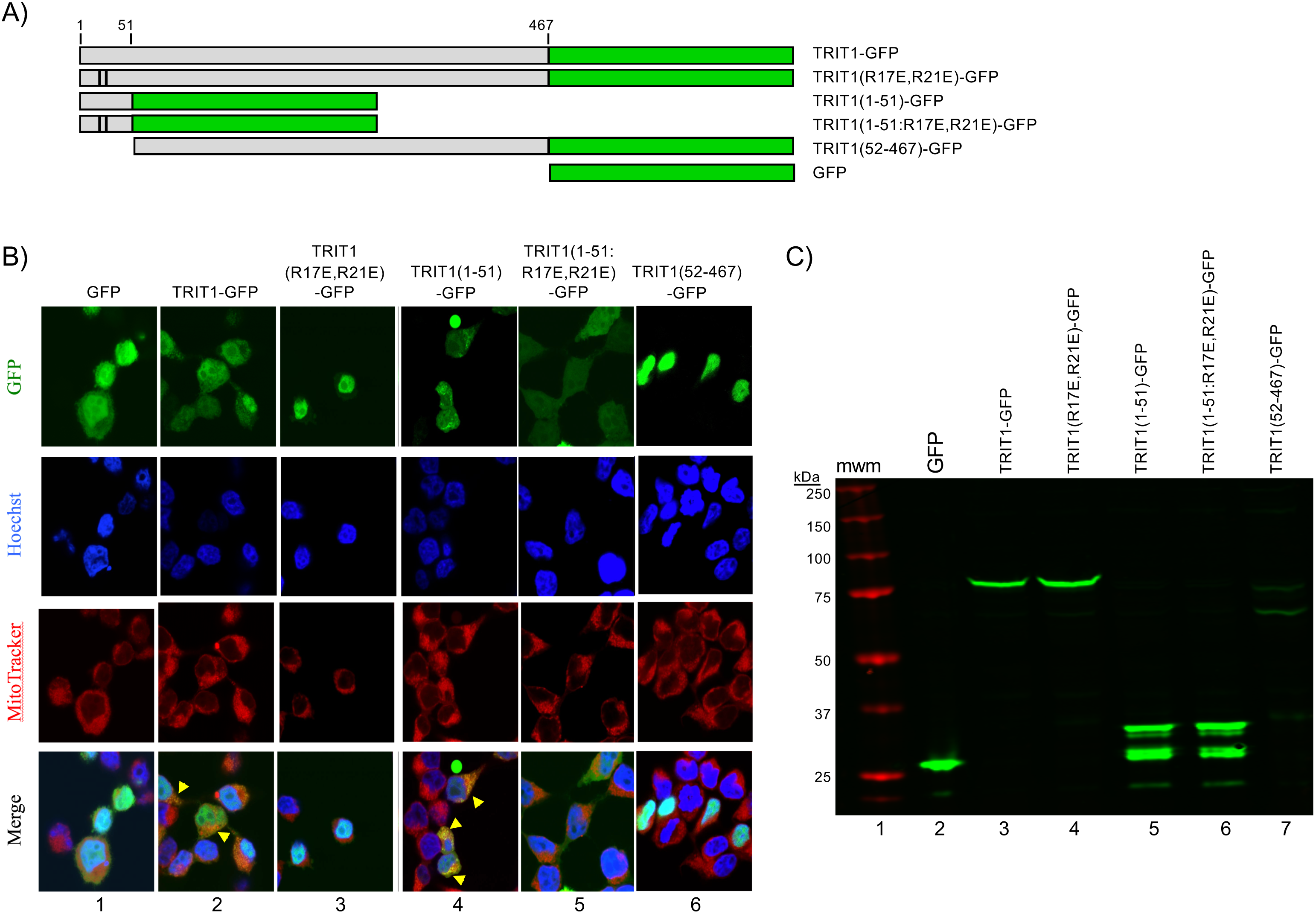
Human TRIT1 contains an N-terminal mitochondrial targeting sequence (MTS). **A)** Cartoon of the GFP-fusion constructs used. Numbering represents amino acids; black stripes indicate positions of two mutated residues in the R17E/R21E constructs. **B)** Representative confocal microscopic images from HEK293 cells transfected 48 hours prior with the constructs in A and stained with Hoechst and MitoTracker. Arrowheads in Merge panels point toward colocalized GFP and MitoTracker. **C)** Western blot from cells transfected with constructs in A) using anti-GFP

The strongest evidence of mitochondrial localization by the TRIT1 MTS came from TRIT1(1-51)-GFP. First, GFP alone produced a diffuse pattern in nuclei and cytoplasm, with nuclei generally more intense (column 1). By contrast, TRIT1(1-51)-GFP (column 4) was more intense in the cytoplasm and with punctate foci that colocalized with MitoTracker. Second, these characteristics including colocalized punctate foci were not observed with the point mutant, TRIT1(1-51:R17E/R21E)-GFP (Fig 2B, column 5). The data support the predicted amphipathic helix as a part of the TRIT1 MTS because substitution of R17 and R21 impaired mitochondrial targeting.

Data here and not shown provide evidence that TRIT1 contains multiple trafficking elements that distribute it to different subcellular compartments, similar to but distinct from Mod5 [45-47, 49, 59, 60]. Mutation of the MTS shifts distribution away from mitochondria. Importantly, as will be shown in a later section, the R17E/R21E MTS mutations specifically impair i6A37 modification of mt-tRNA, but not cy-tRNA modification by TRIT1.

### Complementation of cy-tRNA^Ser^-mediated suppression in *S. pombe*

It was shown by monitoring the codon-specific insertion of an amino acid in the active site of β-galactosidase that is critical for enzymatic activity, that i6A37 increases the decoding activity of cy-tRNA^**Cys**^GCA by ∼3-fold in *S. pombe* [42]. Here, we used a codon-specific reporter for cy-tRNA^**Ser**^UCA-TMS, to examine if TRIT1 mutants can functionally complement *S. pombe* deleted of its IPTase, *tit1Δ* (Fig 3A). In this TMS assay, isopentenylation of cy-tRNA^**Ser**^UCA is required for suppression of premature translation termination at a UGA stop at codon position 215 of *ade6-704* mRNA [61], which results in a decrease in intracellular red pigment [32]. The control strain yYH1 (*tit1*^***+***^) used as a reference, and the test strain yNB5 (*tit1Δ*), contain the same suppressor-tRNA^**Ser**^UCA allele [17]. yYH1 (*tit1*^***+***^) is pink (moderately suppressed) whereas yNB5 (*tit1Δ*) transformed with empty vector pRep82X is unsuppressed, red (Fig 3A, sectors 1 and 2). Transformation of yNB5 with pRep-Tit1 under transcriptional control of the *tit1*^***+***^ promoter, or pRep82X-TRIT1 under the *nmt1*^***+***^ promoter led to suppressed, lighter color yeast (sectors 3 and 4). We note that pRep82X carries the weakest *nmt1*^***+***^ promoter available in the pRepX expression vectors [62, 63]. pRep-Tit1-T12A is inactive [17] (sector 5). pRep82X-TRIT1-T32A and pRep82X-TRIT1(57-467) are also inactive (sectors 6 and 8). pRep82X-TRIT1(R17E/R21E) led to suppression (sector 7) comparable to TRIT1 (sector 4), providing evidence that the mutations that interfere with mitochondrial targeting in human cells do not impair cy-tRNA^**Ser**^UCA-TMS in *S. pombe*.

**Figure 3.**
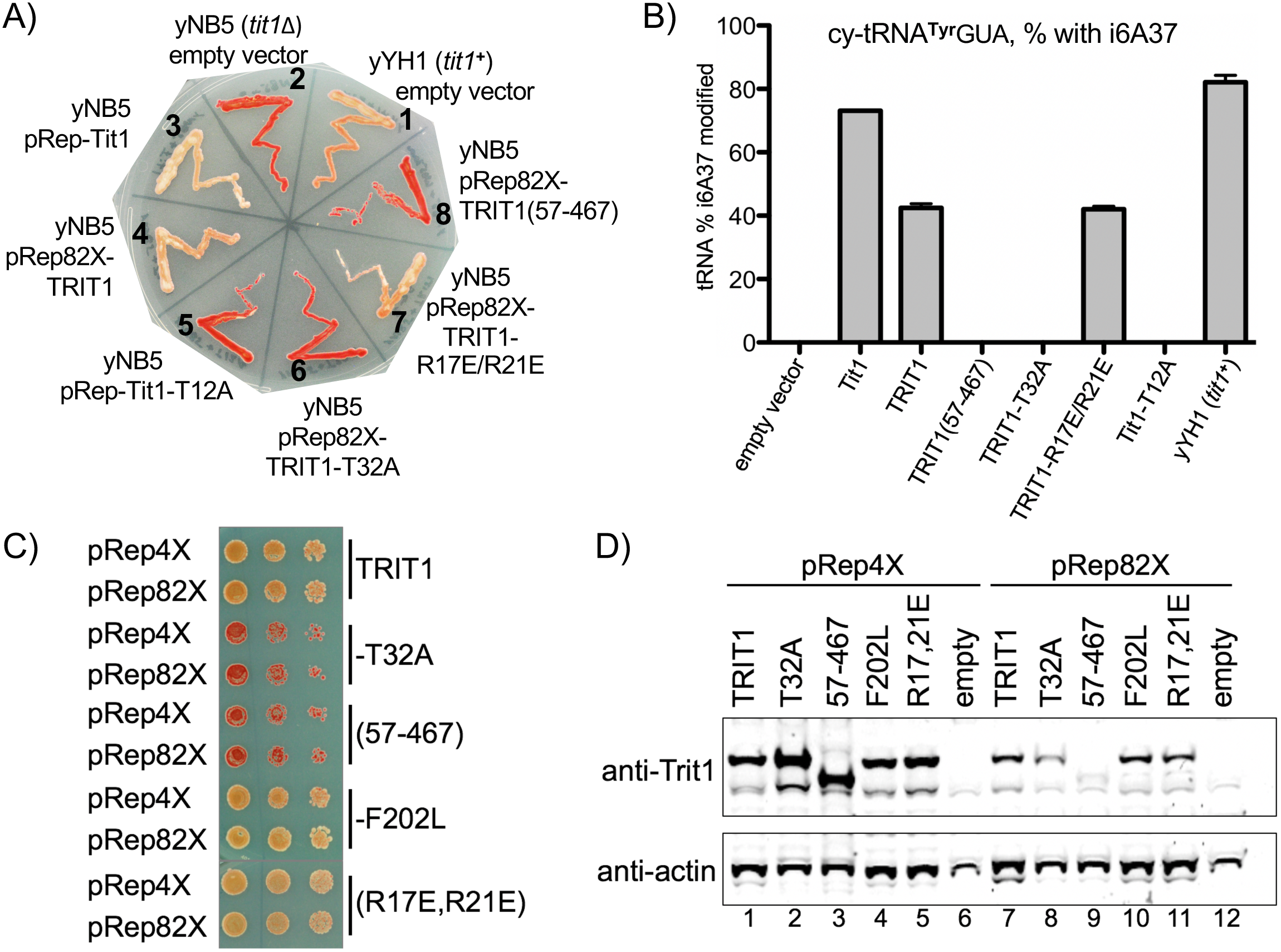
Critical determinants of TRIT1 activity for cy-tRNA substrates. **A)**cy-tRNA^**Ser**^UCA-mediated suppression (TMS) assay of *S. pombe* IPTase-deleted strain yNB5 (*tit1Δ*) transformed with empty plasmid pRep82X or plasmids expressing the proteins indicated (see text), and the *tit1*^***+***^ control strain, yYH1. **B)** Results of PHA6 assay (Methods) that shows relative i6A37 modification efficiency for *S. pombe* endogenous cy-tRNA^**Tyr**^GUA in transformed strains as indicated, calculated as described in methods; error bars represent the spread in two replicates. **C)** TMS assay as in A but displayed as dilution spot assay, for plasmids expressing TRIT1 and variants from the weak (pRep82X) or strong (pRep4X) *nmt1*^***+***^ promoter as indicated on the left. **D)** Western blot of extracts of the cells in C) detected with anti-Trit1 antibody (top panel), and anti-actin antibody (bottom).

RNA from cells represented in Fig 3A were examined for modification of cy-tRNA^**Tyr**^GUA as determined by an i6A37-sensitive northern blot assay (methods, Fig 3B). This confirmed that TRIT1-T32A, TRIT1(57-467) and Tit1-T12A were inactive while TRIT1 and TRIT1(R17E/R21E) were active for i6A37 modification, and Tit1 was more active (Fig 3B).

### Moderate over-expression does not rescue inactive TRIT1 alleles for cy-tRNA^Ser^UCA TMS

As noted in the Introduction, a TRIT1(57-467) isoform predicted to encode a protein beginning with MQVYEGLD (see Fig 1) was active for cy-tRNA^**Tyr**^**UAA**-TMS in *S. cerevisiae* [33]. Because pRep82X-TRIT1(57-467) was inactive for cy-tRNA^**Ser**^-TMS in *S. pombe* (Fig 3), we suspected the discrepancy between the studies might be due to expression levels; use in *S. cerevisiae* of a high copy plasmid with a very strong promoter (*GAL1*) would produce large amounts of TRIT1(57-467) [33]. We therefore compared TMS activity from expression constructs with the weakest and strongest *nmt1*^***+***^ promoters, pRep82X and pRep4X, respectively (Fig 3C). These *nmt1*^***+***^ promoters fused to chloramphenicol acetyltransferase (CAT) or *LacZ*, showed 45-75 fold differences in output [62, 63]. pRep4X-TRIT1(57-467) did indeed produce substantially more accumulation of TRIT1(57-467) than did pRep82X-TRIT1(57-467) (Fig 3D), although without a hint of increased TMS activity relative to pRep82X-TRIT1(57-467) (Fig 3C). Likewise, TRIT1-T32A remained inactive in pRep4X (Fig 3C) as did Tit1-T12A (not shown). TRIT1-F202L, was as active as TRIT1 and also not limited by expression levels, and this was also true for TRIT1-(R17E/R21E) (Fig 3C).

### TRIT1 is severely and specifically deficient for cy-tRNA^Trp^CCA modification

Distinct combinations of tRNAs-i6A37 contribute to the cytosolic translation machineries of *S. cerevisiae, S. pombe* and human cells (Table 1). As noted, cy-tRNA^**Trp**^CCA carries unmodified A37 in *S. cerevisiae* despite an A36-A37-A38 sequence [15], consistent with poor substrate activity for Mod5 IPTase; in contrast, the *S. pombe* IPTase, Tit1 most readily modifies this and all cy-tRNAs^**Trp**^CCA tested [17]. Limited experimental data suggest that some of this discrimination resides in a variable anticodon recognition loop of the IPTase [17] (Discussion). More relevant here is that both the human and *S. cerevisiae* mt-tRNAs^**Trp**^ which contain i6A37 are encoded to produce <u>U</u>CA anticodons in which the U34 obtains a bulky modification, rather than CCA (Table 1, Discussion). Therefore, we wanted to examine the extent to which TRIT1 could modify *S. pombe* cy-tRNA^**Trp**^CCA and mt-tRNA^**Trp**^CCA.

TRIT1 was indeed deficient for modification of cy-tRNA^**Trp**^CCA, reminiscent of Mod5, but more severe when compared side-by-side (Fig 4A-C). Fig 4A shows results of an i6A37-sensitive PHA6 blot assay for cy-tRNAs ^**Trp**^CCA, ^**Tyr**^GUA and ^**Ser**^AGA. The PHA6 (<u>P</u>ositive <u>h</u>ybridization in <u>A</u>bsence of i<u>6</u>A37) assay is based on loss of oligo-DNA probe hybridization to the ACL due to presence of i6A37, and is calibrated for quantitation by a body probe to the T stem loop on the same tRNA (Fig 4B, C) [14, 17, 42, 64]. In addition to Mod5, we included two mutants, Mod5-M1A and Mod5MTS-TRIT1, that are important below for assessing modification of the mt-tRNA^**Trp**^CCA. TRIT1 (and Mod5MTS-TRIT1) is deficient for cy-tRNA^**Trp**^CCA modification as reflected by strong hybridization signal in lane 1 relative to lanes 2 (Mod5) and lane 6 (Tit1) of Fig 4A upper panel. These signal differences are less dramatic for cy-tRNA^**Tyr**^GUA (Fig 4A, middle panel) and cy-tRNA^**Ser**^AGA (lower panel) (Fig 4A). Thus, cy-tRNA^**Trp**^CCA exhibits very low substrate activity specific to TRIT1 (and Mod5MTS-TRIT1), and relative to cy-tRNAs ^**Tyr**^GUA and ^**Ser**^AGA in the same cells. By contrast to TRIT1, cy-tRNA^**Trp**^CCA is most efficiently modified by Tit1 followed by Mod5 (Fig 4A, lanes 6, 2, 1; and compare black with gray bars in Fig 4C). The pattern obtained with ectopic pRep-Tit1 was similar to yYH1(*tit1*^***+***^) (not shown).

**Figure 4.**
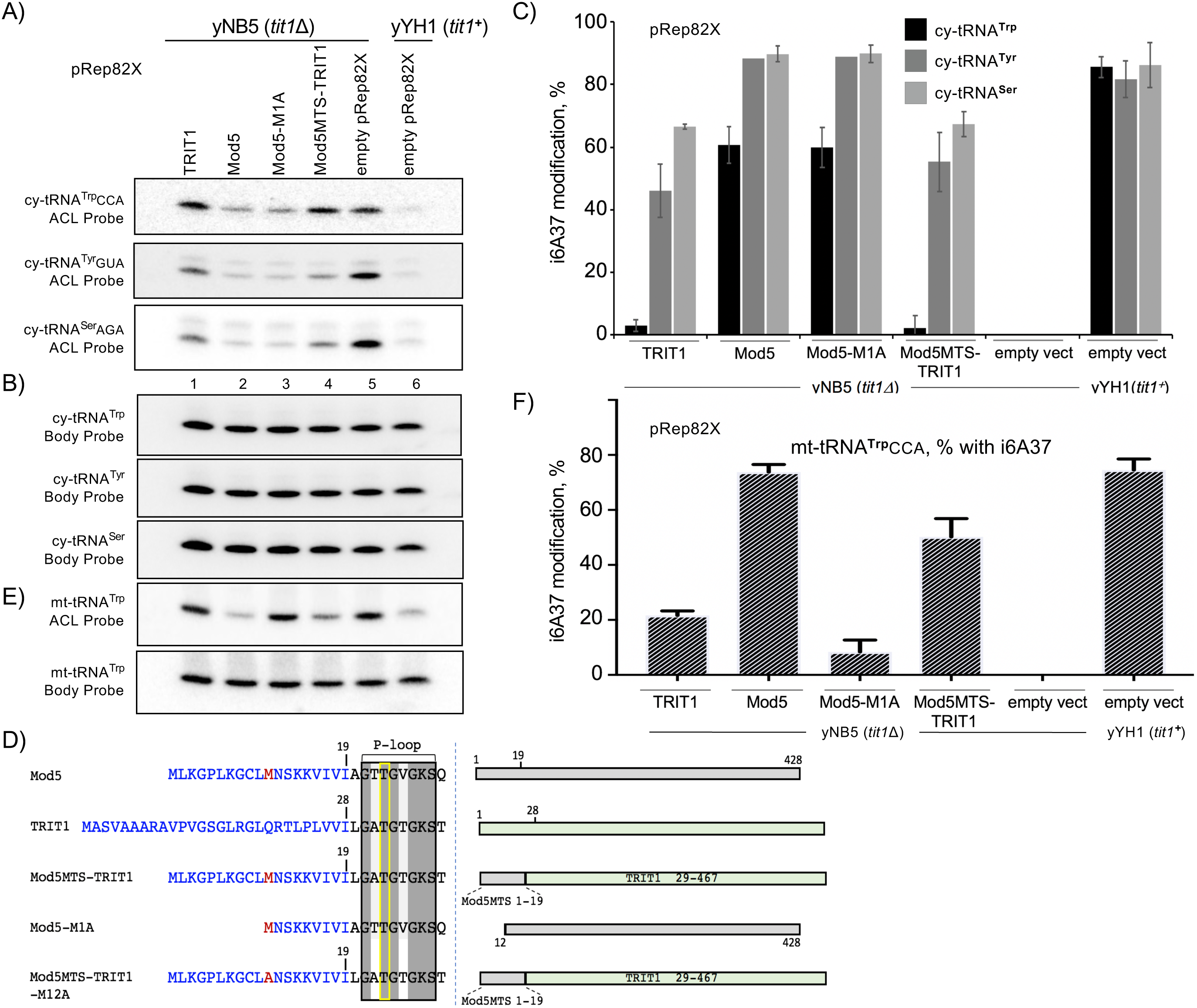
TRIT1 is specifically deficient for cy-tRNA^Trp^CCA modification. **A)** i6A37-sensitive PHA6 northern blot assay for three tRNAs from *S. pombe* cells with pRep82X expression constructs indicated above the lanes. The blot was sequentially probed with i6A37-sensitive ACL probes to the cy-tRNAs indicated to the left of each panel. **B)** The body probes, to the T stem loop, for each of the cy-tRNA shown in A. **C)** Quantitation of % i6A37 modification of the three cy-tRNAs from biological duplicate PHA6 northern blots as for A & B; n=5 for cy-tRNA^**Trp**^CCA and all others except n=1 for cy-tRNA^**Tyr**^ in Mod5 and Mod5M1A. **D)** Schematic showing constructs used for analysis of mt-tRNA^**Trp**^CCA i6A37 modification in *S. pombe*. Left: amino acid sequence depiction of N-termini only, similar to Fig 1; invariant Thr is demarcated by yellow rectangle. Alternative translation initiation sites are shown in red. Right: linear representation of the constructs with landmarks in the N-terminal regions indicated. **E)** The same blot as in panels A-B, probed for mt-tRNA^**Trp**^CCA using ACL and body probes as indicated. **F)** Quantitation of % i6A37 modification of mt-tRNA^**Trp**^CCA from biological duplicate PHA6 northern blots.

### TRIT1 modifies mt-tRNA^Trp^CCA in *S. pombe*

Analysis of *S. pombe* mt-DNA by tRNAscan-SE predicted 25 mt-tRNAs (including 3 for Met), of which two, mt-tRNA^**Trp**^CCA and mt-tRNA^**Cys**^GCA contain an AAA recognition element. Prior studies using standard PHA6 assay conditions accounted for i6A37 modification of mt-tRNA^**Trp**^CCA at 25-30% in yYH1(*tit1*^***+***^) cells but only background levels on mt-tRNA^**Cys**^GCA, while it was >75% on cy-tRNAs on the same blots [14, 42]. For the present study, we determined that increasing the post-hybridization wash temperature, to 10-12.5°C above standard conditions is necessary to reveal a difference in signal between IPTase positive and negative sample mt-tRNAs (see Supp Fig S1). As shown below, these optimal conditions revealed mt-tRNA^**Trp**^CCA modification at ∼75% for Tit1 and Mod5 but reproducibly only ∼20% for TRIT1 (Fig 4D-F). By contrast, cy-tRNA^**Ser**^AGA was modified to ∼70% by TRIT1 in the same cells, and 80-90% by Mod5 and Tit1 (Fig 4A-C).

We considered that multiple variables may contribute to tRNA i6A37 modification efficiency. In the cytosol, cy-tRNA^**Trp**^CCA is an inherently poor substrate that must compete for IPTase activity with variable-abundance multi-copy tRNA gene-derived cy-tRNA^**Tyr**^GUA, cy-tRNA^**Ser**^AGA, and cy-tRNA^**Ser**^UGA. In mitochondria, competition is predictably less because mt-tRNA^**Trp**^CCA and mt-tRNA^**Cys**^GCA sequences are single copy in mt-DNA. Another variable for mt-tRNAs may be the IPTase concentration in the mitochondrial matrix, which is likely governed by their MTS activity.

It was previously shown that the MOD5 MTS is required for i6A37 modification of mt-tRNA^**Trp**^CCA in *S. pombe* [14]. We examined MTS activity by first comparing N-terminal fusions as schematized in Fig 4D; codons 1-28 of TRIT1 were replaced with codons 1-19 of MOD5. We confirmed that Mod5-M1A, which initiates at M12 and lacks a MTS [45, 46], was deficient for mt-tRNA^**Trp**^CCA modification relative to Mod5 (Fig 4E, F). Fig 4E, F also show that Mod5MTS-TRIT1 exhibited >2-fold more modification of mt-tRNA^**Trp**^CCA than TRIT1. In any case, TRIT1 and Mod5MTS-TRIT1 led to higher levels of modification of mt-tRNA^**Trp**^CCA than of cy-tRNA^**Trp**^CCA (Fig 4C vs. 4F, and below). These results suggest that TRIT1 can modify mt-tRNA^Trp^CCA more efficiently than cy-tRNA^Trp^CCA in *S. pombe* as will be corroborated in Fig 6 below.

**Figure 5.**
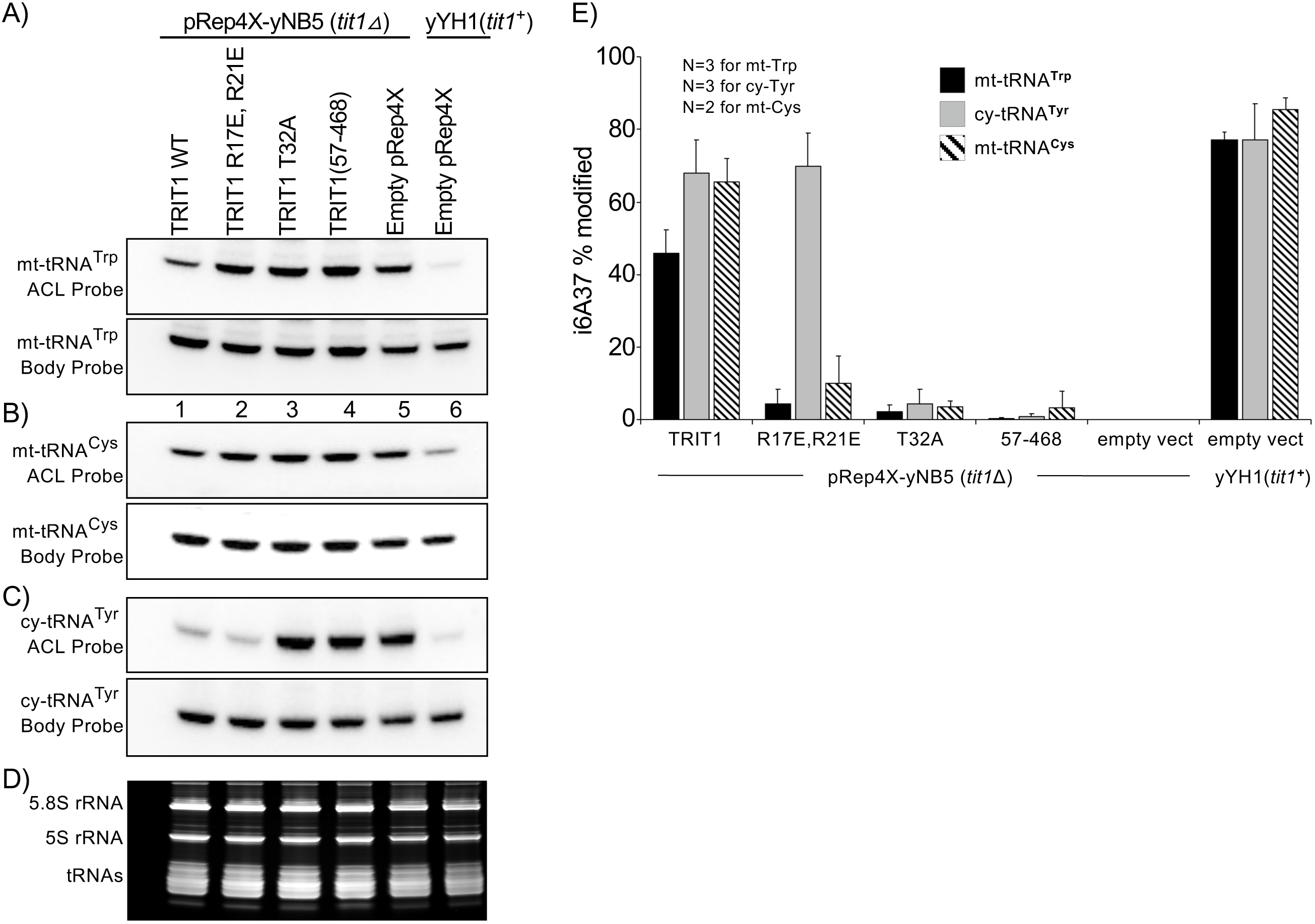
TRIT1 MTS-dependent i6A37 modification of mt-tRNA^Trp^CCA and mt-tRNA^Cys^GCA. **A-C)** Northern blot panels probed for mt-tRNA^Trp^CCA, mt-tRNA^Cys^GCA and cy-tRNA^Tyr^GUA with ACL and body probes as indicated; and **D)** EtBr panel. **E)** Quantitation of % i6A37 modification of the mt-tRNAs and the cy-tRNA from biological replicate PHA6 northern blots as indicated.

**Figure 6.**
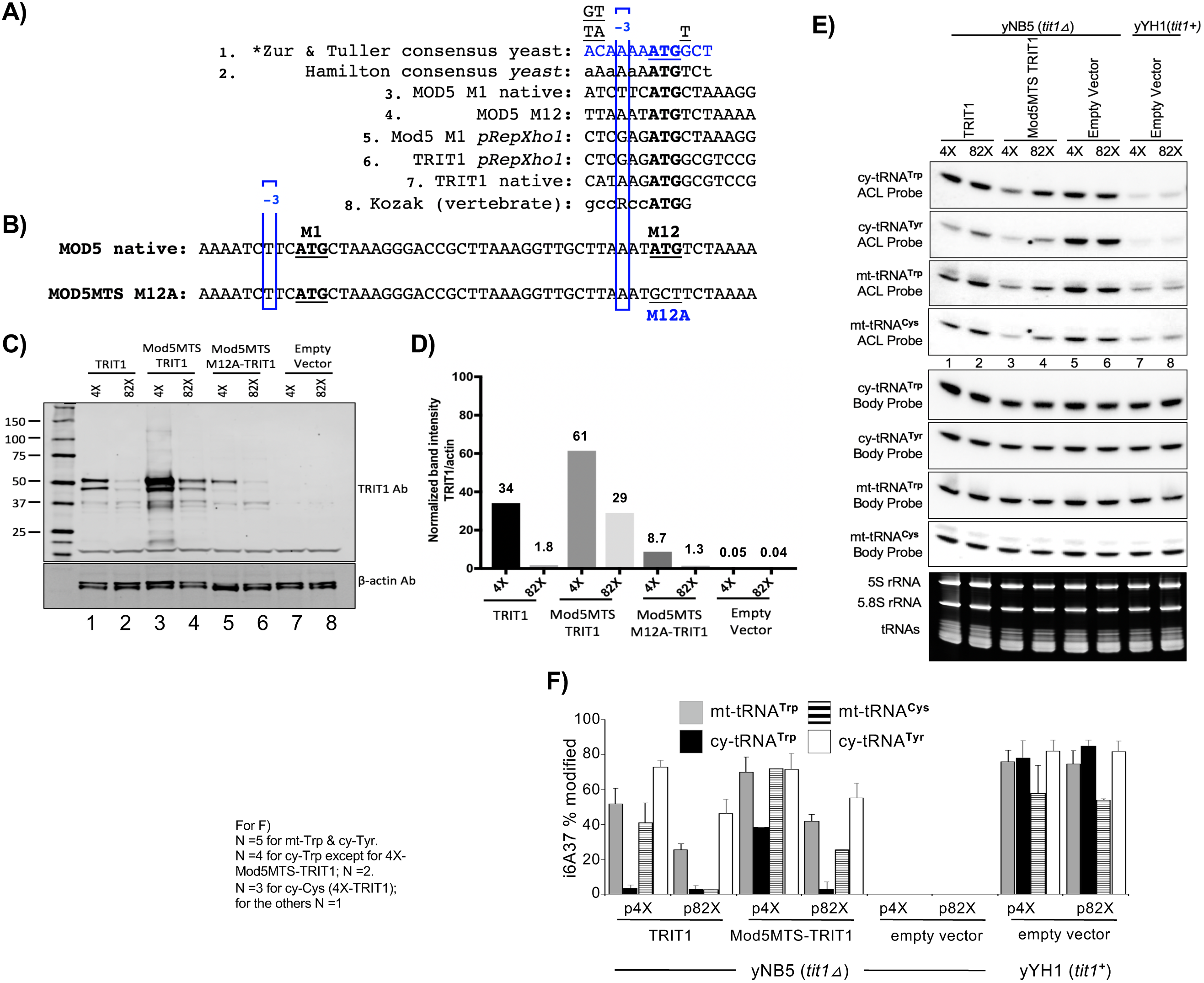
Concentration-dependent i6A37 modification of cy-tRNA^Trp^CCA. **A)** Translation initiation contexts of ATG codons; the first two lines show consensuses derived for *S. cerevisiae* [highly expressed genes 67, 89] and the last line shows a consensus derived for vertebrate [90]. Lines 3 and 4 are for the alternative ATGs that encode MOD5 M1 and M12 in their native context; lines 5 and 6 are for the MOD5 M1 and TRIT1 M1 after cloning into the *Xho1* site of *S. pombe* expression vectors pRep4X or pRep82X (*pRepXho1*). Line 7 shows the native TRIT1 M1 ATG context. **B)** Nucleotide sequence of MOD5 including its M1 and M12 ATG sites and the M12A substitution mutation in the Mod5MTS-M12A-TRIT1 construct. **C)** Western blot of TRIT1 protein developed using anti-TRIT1 Ab, from cells with constructs indicated above the lanes containing the strong (pRep4X) or weak (pRep82X) *nmt1*^***+***^ promoter (see text). Lanes are numbered below. MW markers are indicated in kDa. The common band below 25 kDa is an internal control. Lower panel shows β-actin as loading control used for quantitative normalization. **D)** Determination of TRIT1 levels in C by quantitative Odyssey CLx imaging (Methods); numbers above bars indicate TRIT1/β-actin levels in each sample. **E)** Northern blot of 2 cy- and 2 mt-tRNAs by TRIT1 and Mod5MTS-TRIT1 each from the strong promoter, pRep4X and weak promoter pRep82X, as indicated above the lanes. The top four panels show the ACL probings as indicated to the left, and the bottom four panels show the corresponding body probings. **F)** Quantitation of % i6A37 modification of the mt-tRNAs and the cy-tRNAs as indicated. **G)** Clover leaf representations of *S. pombe* cy-tRNA^**Trp**^CCA and mt-tRNA^**Trp**^CCA as encoded by the nuclear and mitochondrial DNA and folded by tRNAscan-SE [81] (Table 1, Discussion).

### TRIT1 MTS-dependent i6A37 modification of mt-tRNA^Trp^CCA and mt-tRNA^Cys^GCA

We functionally validated the TRIT1 MTS in *S. pomb*e by examining mt-tRNA modification (Fig 5A-E). Raising TRIT1 levels with the strongest *nmt1*^***+***^ promoter, pRep4X-TRIT1 increased mt-tRNA^**Trp**^CCA modification to ∼45% (Fig 5A, E), higher than the ∼20% achieved with pRep82X-TRIT1 (Fig 4F). Importantly, biological triplicate determinations revealed that the MTS mutations in pRep4X-TRIT1(R17E/R21E) lowered mt-tRNA^**Trp**^CCA modification to ∼6% while modification of cy-tRNA^**Tyr**^GUA was not affected by these mutations, and the negative control proteins, TRIT1-T32A and TRIT1(57-467) showed negligible modification (Fig 5A, E). Similar to mt-tRNA^**Trp**^CCA, mt-tRNA^**Cys**^GCA modification was negligible with pRep4X-TRIT1(R17E/R21E) (Fig 5B, E). We note that mt-tRNA^**Trp**^CCA and mt-tRNA^**Cys**^GCA were determined to be 80% and 90% modified respectively in yYH1(*tit1*^***+***^) cells (Fig 5B, E). Thus, the R17E/R21E mutations that impaired TRIT1 MTS-mediated localization in HEK293 cells, also impaired modification of mt-tRNA^**Trp**^CCA and mt-tRNA^**Cys**^GCA in *S. pombe* mitochondria, while they had no effect on cy-tRNA^**Tyr**^GUA (Fig 5E) nor on cy-tRNA^**Ser**^UCA-mediated TMS (Fig 3A). The data provide strong evidence that the TRIT1 MTS is functional in *S. pombe.*

### TRIT1 concentration-dependent modification of the inferior substrate, cy-tRNA^Trp^CCA

We wanted to examine if further elevation of cytosolic TRIT1 levels could lead to increased modification of cy-tRNA^**Trp**^CCA. For this we made use of the Mod5MTS sequence for its high level production of cytosolic protein due to the very favorable context of its M12 ATG for translation initiation [65], for TRIT1 expression in *S. pombe* (Fig 6A-C). Studies of MOD5 have shown that suboptimal context of its first ATG (M1) accounts for relatively low levels (∼10%) of the MTS-containing isoform, IPPT-I, as compared to the IPPT-II isoform (∼90%) that begins at ATG (M12), which resides within a strong consensus for efficient initiation [65-67] (Fig 6A,B). As cloned in the *Xho*I sites of pRep 82X and 4X, the context upstream of ATG (M1) of both TRIT1 and Mod5MTS-TRIT1 constructs are identical but suboptimal because they lack A at −3, a most conserved and influential position [66, 67] (Fig 6A, *pRepXho1*). This suggests that both pRepX-TRIT1 and pRepX-Mod5MTS-TRIT1 undergo comparable ‘leaky scanning’ (i.e., inefficient) initiation at their first ATG, M1, and that a higher level of TRIT1 produced by Mod5MTS-TRIT1 would be due to more efficient initiation at the second ATG, M12, the cytoplasmic isoform, as a result of its better context (Fig 6A, B). Examination of TRIT1 expression was largely consistent with these expectations (Fig 6C, D, the calculated mass of full length TRIT1 is 52.7 kDa). As expected, the Mod5MTS-TRIT1 constructs produced higher levels than the corresponding pRep4X and pRep82X constructs (Fig 6C, lanes 1-4). Importantly, blocking initiation of the cytoplasmic-specific isoform with the substitution mutation, M12A, reduced TRIT1 protein levels by ∼10-fold (Fig 6C, D, compare lanes 5 and 3), providing evidence that the excess TRIT1 produced by pRepX-Mod5MTS-TRIT1 over that produced by pRepX-TRIT1 represents cytoplasmic protein.

Indeed, cells expressing higher levels of cytoplasmic TRIT1 exhibited >7-fold increase in modification of cy-tRNA^**Trp**^CCA compared to cells with lower levels (Fig 6E, F, compare black bars for pRep4X-TRIT1 vs pRep4X-Mod5MTS-TRIT1). The cytoplasmic concentration-dependent increase in modification appeared specific for cy-tRNA^**Trp**^CCA which increased >7-fold whereas cy-tRNA^**Tyr**^GUA was unaffected (Fig 6F).

Because TRIT1 does not appear to use alternative translation initiation to control its IPTase distribution to mitochondria and other subcellular locations (see Fig 1, 2), we cannot estimate what fraction of the total TRIT1 protein in lane 1 of Fig 6C is cytoplasmic and/or mitochondrial. To determine the relative concentrations in cytoplasm vs. mitochondrial matrix in human cells is beyond the scope of this study. Examination of the ATG context of native TRIT1 (Fig 6A, line 7) indicates that it resides in a reasonably good Kozak consensus that would direct a large fraction of TRIT1 to the mitochondria, consistent with our GFP-fusion imaging.

## DISCUSSION

In this study we demonstrated that human TRIT1 uses an N-terminal MTS for mitochondrial localization in human cells and also showed it to be functional in *S. pombe*. As reviewed in the Introduction, proper subcellular localization is critical for IPTase function; in *S. cerevisiae*, the mechanisms by which Mod5 localizes to nuclei, cytoplasm and mitochondria have been extensively investigated [45-47, 65, 68, 69]. *S. cerevisiae* Mod5 (as well as *C. elegans* GRO-1) uses alternative translation initiation sites to target to mitochondria or cytoplasm [45, 46, 53]. Our analyses indicate that TRIT1 uses a single translation start site. Mitochondrial activity of TRIT1 is relevant because pathogenic mutations to it that decrease i6A37 modification of both cy-tRNAs and mt-tRNAs, manifest in patients principally as mitochondrial insufficiency due to impairment of the mitochondrial translational system [41] [also see 54]. We should note that the majority of pathogenic SNPs in TRIT1 described [41, 54], do not reside within or near the MTS examined here, (one allele of a compound heterozygous genotype predicts a truncation at Arg-8) [54]. Examining the N-terminal MTS was also important because a popular algorithm predicts presequence cleavage at position 47 which would remove residues critical for modification activity, including the conserved P-loop containing the invariant catalytic T32, which we showed is required for efficient modification activity of cy- and mt-tRNAs. A resolution of this problem was provided by the MitoFates algorithm [56], which is an improved method for predicting cleavable N-termini MTSs, and indeed predicts a two component N-terminal MTS for TRIT1 (Fig 1) but with very low probability of cleavage. Subcellular fractionation had shown that TRIT1 colocalized to the lysate, mitochondria, and the mitochondrial matrix fractions but with no evidence of cleavage [41]. Thus, our data and other analysis favor a model in which the N-terminal MTS directs mitochondrial import of TRIT1 but do not support cleavage. Furthermore, this work has established a *S. pombe* system that can be used to study the efficacy of TRIT1 MTS for functional mt-tRNA i6A37 modification.

Another conclusion is that the TRIT1 IPTase exhibits substrate-specific deficiency for cy-tRNA^**Trp**^CCA modification relative to the Mod5 and Tit1 IPTases, despite the presence of the A36-A37-A38 sequence identity element in this tRNA. The AAA sequence within the anticodon loop was known to be a major identity element for *E. coli* IPTase, and to be influenced by specific base pairs in the anticodon stem, as well as certain characteristics of the loop itself [70]. Our data are consistent with the earlier findings but are distinct in that they show significant differences among eukaryotic IPTases that are relevant to the variable identities of tRNAs-i6A37 found in nature (Table 1). We show that increasing the cytoplasmic concentration of TRIT1 is able to partially overcome the poor substrate activity of the cy-tRNA^**Trp**^CCA substrate (Fig 6).

### The N-terminal MTS of TRIT1

As noted above, MitoProt II [55] predicted an N-terminal MTS for TRIT1 with high probability, and with a post-import cleavage at position 47 which is at odds with our mutagenic analysis and prior characterization of critical catalytic residues in other IPTases, and supported by crystallographic studies in which Mod5 was observed bound to tRNA^**Cys**^GCA together with a DMAPP analog substrate. TRIT1-T32A and Tit1-T12A are homologous to *E. coli* IPTase MiaA-T19A, a substitution that resulted in the largest decrease in *k*_*cat*_ (600-fold) of all mutations tested [17, 42, 43]. The corresponding T23 in the Mod5 structure is positioned adjacent to DMAPP for catalysis [44]. Thus, it is important to emphasize that MitoFates [56], which was designed as an improved prediction method for N-terminal MTSs and their cleavage sites, predicted a N-terminal MTS comprised of a TOM20 receptor binding site followed by an amphipathic helix (Fig 1) but with low probability of cleavage, 0.106, well below the default cutoff. The conclusion that the N-terminal 11 amino acids of Mod5 are necessary for mitochondrial import and that the MTS presequence is probably not proteolytically removed after import [45], are supported by a MitoFates analysis that indeed predicts positively charged amphiphilicity within the first 11 amino acids but with a very low probability of cleavage, of 0.026. That study also indicates that the Mod5 MTS is not fully efficient at import, as a functional pool of N-terminal extended protein remained in the cytoplasm [45]. Our data with Mod5MTS-M12A-TRIT1, are consistent with this as it retained substantial i6A37 modification activity for cy-tRNAs (data not shown).

TRIT1 localizes to the mitochondrial matrix, where modified mt-tRNAs function in human cells, although with no evidence of cleavage, whereas GDH (glutamate dehydrogenase), a matrix-specific protein, exhibited evidence of cleavage [41]. Point mutagenesis of the two key basic residues in the TRIT1 MTS amphipathic helix debilitated mitochondrial localization in human cells (Fig 2) and also debilitated TRIT1 for i6A37 modification of mt-tRNAs, but not cy-tRNAs in *S. pombe* cells (Fig 5). The data indicate that TRIT1 has a N-terminal MTS comprised of a TOM20 receptor binding site followed by an amphipathic helix (Fig 1) but is probably not cleaved.

Amounts of TRIT1 in mitochondrial and submitochondrial fractions, and sensitivity to proteinase treatment relative to GDH raised the possibility that a significant fraction of cytoplasmic TRIT1 may be associated with the outer mitochondrial membrane [41]. We note that such a possibility is consistent with our GFP-fusion construct imaging in which a substantial amount of cytoplasmic TRIT1(1-51)-GFP and TRIT1-GFP would appear to be associated with mitochondria (Fig 2B).

### IPTase function and the variable identities of their associated tRNAs-i6A37

Data reported here support the idea that IPTases can discriminate against substrate tRNAs in an anticodon identity type manner, at least with regard to the TrpCCA anticodon, consistent with Mod5 structural studies that show base-specific contacts to G34 of tRNA^**Cys**^ [44]. Biochemical and *in vivo* data showed that a point substitution to the anticodon of cy-tRNA^**Trp**^CCA from C34 to G34 rescued its substrate activity for Mod5 [17]. A question that arises is how might this relate if at all, to the variable identities of tRNAs-i6A37 that are found in different eukaryotes?

The i6A37 modification was known to increase ribosome binding [71] and functional efficiency of several suppressor-tRNAs derived from natural tRNAs that decode UNN codons, although to different extents [72]. Early work led to the proposal that i6A37 could stabilize the otherwise weak base pairing between the codon U1 base and anticodon base A36, and structural evidence supports this for tRNA^**Phe**^GAA-ms2i6A37 bound to a UUC codon in the decoding center of the ribosome [73]. Studies in *S. pombe* supported the conclusion that i6A37 increases the translational activity of tRNAs by 3-to-4 fold generally [42]. This is consistent with the idea that i6A37 increases the affinity for ribosome binding and the functional efficiency for translation of tRNA [71]. Recent high resolution data from *mod5-*deletion cells that revealed dramatic decreases in A site ribosome occupancies over all of the *S. cerevisiae* tRNA-i6A37-cognate codons [74], fit with a model in which absence of i6A37 decreases the affinity of the associated tRNAs for the translating ribosome.

However, apart from increasing the general translation activity of associated tRNAs, i6A37 may exert other, specific activities, for example, protection against frameshifting and mistranslation by near-cognate tRNAs, in a codon-specific and/or context-specific manner [42, 75, 76]. Lines of evidence indicate that i6A37 and/or its derivatives can exhibit such tRNA-specific and codon- and/or context-dependent effects [see 72]. We should therefore suspect that some of the variability of tRNA-i6A37 identities may reflect that in eukaryotes, in which tRNA gene copy number is variable and dynamic [77-81], other mechanisms may compensate for lack of i6A37 on some tRNAs.

Differences in the identities of tRNAs that have been selected or not for i6A37 modification can be due to sequence differences in the 37 and 38 positions, which contribute to the major IPTase identity element, A36-A37-A38 or to species-specific characteristics of the IPTase, the latter as first reported for Mod5 [17] and extended in this study to TRIT1. Thus, cumulative data now more firmly indicate that both mechanisms contribute to tRNA-specific selection for i6A37 modification in the cytosolic and mitochondrial translation systems of eukaryotes (Table 1), and that this identity appears to reside at least to some extent, i.e., for cy-tRNA^**Trp**^CCA, in the anticodon itself.

The i6A37 modification is found on nine of ten bacterial tRNAs that bear NNA anticodons (Table 1). This suggests that ancient IPTases had broad recognition accommodation of tRNAs with UNN codons [70] with the exception of tRNA^**Ser**^GGA even though it bears A36-A37-A38, although this is not a substrate in eukaryotes which do not have functional genes for this tRNA [16, 81-83]. The data that show that both Mod5 and TRIT1 exhibit low activity for cy-tRNA^**Trp**^CCA relative to Tit1 and to other cy-tRNAs with the A36-A37-A38 identity element is consistent with the idea that eukaryotic IPTases and their translation systems evolved decreased requirements among the UNN decoding tRNAs for i6A37 modification (Table 1).

### Insight into discrimination against CCA anticodon recognition by eukaryotic IPTases

It was previously reported that budding yeast IPTase, Mod5 discriminates against the cy-tRNA^**Trp**^CCA substrate whereas fission yeast Tit1 does not [17]. It is clear from the present work that mammalian TRIT1 IPTase also discriminates against the cy-tRNA^**Trp**^CCA substrate. Although humans have a naturally limited cy-tRNA substrate repertoire [64] (Table 1), examining TRIT1 in *S. pombe* along with Mod5 afforded us the opportunity to assess its activity on an array of substrates not otherwise readily available in an *in vivo* setting. This revealed two interesting results, that MOD5 exhibited more activity for *S. pombe* cy-tRNA^**Trp**^CCA in *S. pombe* than it exhibits for *S. cerevisiae* cy-tRNA^**Trp**^CCA in *S. cerevisiae*, and that TRIT1 was more severely impaired than Mod5 for modification of *S. pombe* cy-tRNA^**Trp**^CCA. With regard to the former we note that Mod5 expression in *S. pombe* was directed by the strong *nmt1*^***+***^ promoter from the high copy plasmid (∼20/cell). Because accumulation *in vivo* can be influenced by protein sequence as well as mRNA determinants, comparison of different IPTases at the same concentration is not expected. Nonetheless, the low activity of *S. pombe* cy-tRNA^**Trp**^CCA as a Mod5 substrate as compared to cy-tRNA^**Tyr**^GUA, cy-tRNA^**Ser**^UCA and cy-tRNA^**Ser**^AGA, and as compared to Tit1 which modifies cy-tRNA^**Trp**^CCA as well as the other substrates, was evident (Figs 3A, 4C). It was further evident that TRIT1 was more severely specifically impaired for cy-tRNA^**Trp**^CCA modification than Mod5 (Fig 4C).

An unexplained observation is the ∼10-fold disparity with which pRep4X-TRIT1 modifies the mitochondrial and cytoplasmic tRNAs ^**Trp**^CCA, at 50% and 4%, respectively in *S. pombe* (Fig 6F). Although the basis for this is unknown, as described below, we suspect differential position 34 modification. This in part because the DNA encoding the anticodon loops and the proximal base pair at the stem are identical in these tRNAs, and each of the other stem-loops share the same number of nucleotides (Fig 7A-C). As noted, the exact sequence of the anticodon stem of the poor substrate, tRNA^**Ser**^GGA significantly impacts activity for the MiaA IPTase [70]. However, this system of nucleotide identity that is applicable to MiaA:tRNA^**Ser**^GGA anticodon stem recognition cannot account for the disparity here since the most negative element in that study, a G30-U40 base pair, is present in the more favorable TRIT1 substrate, mt-tRNA^**Trp**^CCA but replaced by G30-C40 in the less favored cy-tRNA^**Trp**^CCA, and the second most positive feature, purines at positions 29 and 30, is lacking in mt-tRNA^**Trp**^CCA but present in cy-tRNA^**Trp**^CCA (Fig 7A, B). Thus, either TRIT1 does exhibit anticodon stem recognition but is very different from that of MiaA or another basis explains the apparent discrimination by TRIT1 observed for cy-relative to the mt-tRNA^**Trp**^CCA.

**Figure 7.**
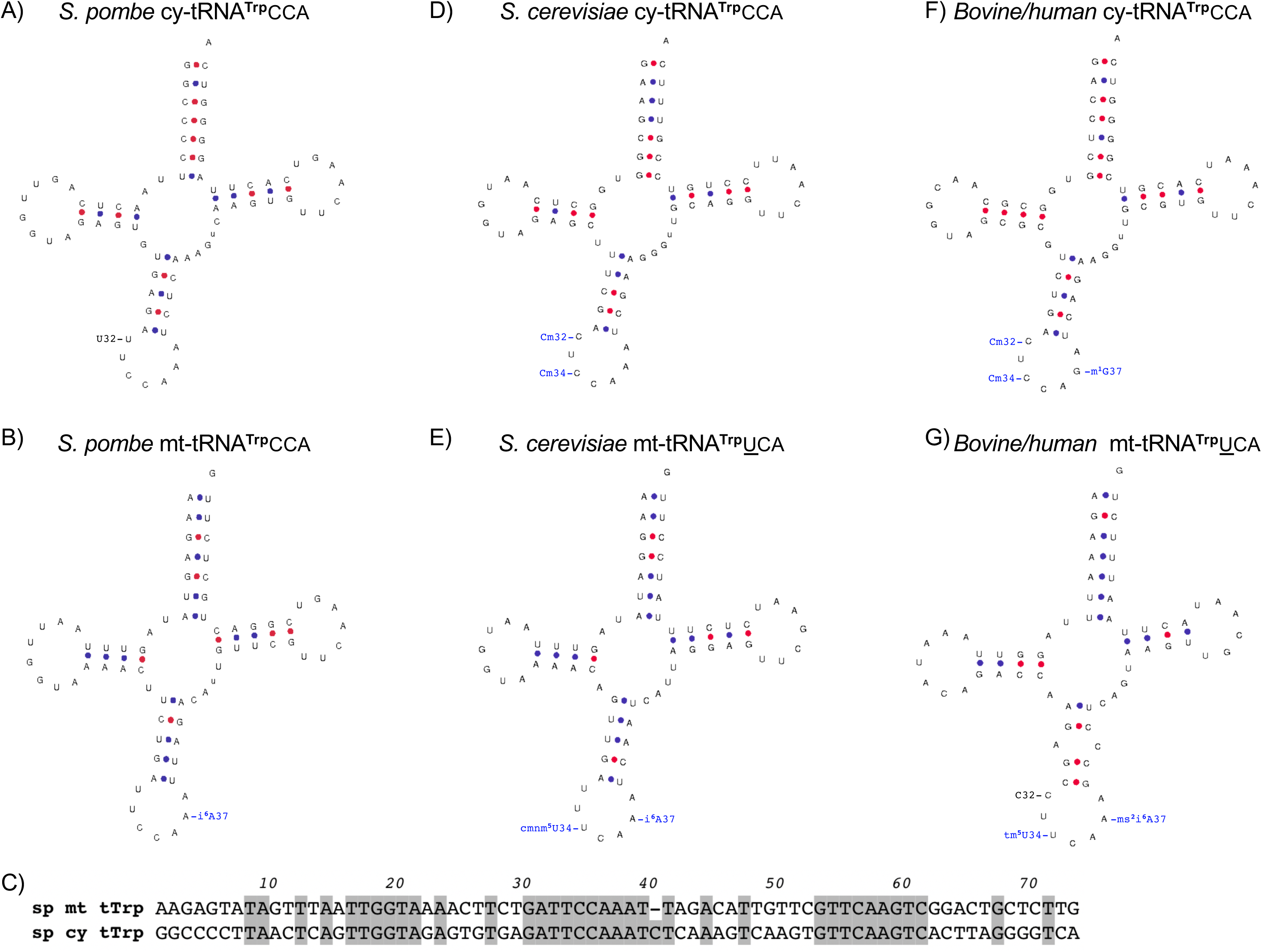
Secondary structures of the cy- and mt-tRNAs^Trp^ as predicted by the tRNAscan-SE-folding algorithm for *S. pombe* (A, B), *S. cerevisiae* (D, E) and Bovine/Human (F, G). Known modifications, to the anticodon loop only, are annotated as blue text, as indicated; black annotated text indicates encoded nucleotide identity that differs among species [12] [15]. **C)** Sequence alignment of *S. pombe* mt- and cy-tRNAs^Trp^.

Because the mt- and cy-tRNAs^**Trp**^CCA exhibit 10-fold difference in substrate activity for TRIT1 in the cytosol and mitochondria of *S. pombe*, but are identical in their anticodon loops and of similar structure elsewhere with ∼50% sequence identity (Fig 7C), based on available data, it should be considered that they may be differentially modified on C34. First, the C at position 34 most strongly sensitizes Mod5 to the negative influence of the CCA anticodon [17].

Although the modification status of *S. pombe* mt- and cy-tRNAs^**Trp**^CCA are unknown, *S. cerevisiae* cy-tRNA^**Trp**^CCA carries Cm at 34 whereas *E. coli* and some other prokaryotes carry unmodified C34 [84]. As noted (Table 1), mammalian and *S. cerevisiae* mt-tRNA^**Trp**^UCA are encoded with U at position 34 and are modified with the bulky groups, tm^**5**^U and cmnm^**5**^U, bearing cmnm U34 and A36-respectively (Table 1) [12, 84, 85]. Because Mod5 doesn’t modify *S. cerevisiae* cy-tRNA^**Trp**^CCA bearing Cm34 and A36-A37-A38, but does modify mt-tRNA^**Trp**^UCA ^**5**^A37-A38, it is tempting to speculate that the bulky modified uridines specifically enhance the activities of their substrate tRNAs for the Mod5 and TRIT1 IPTases. Data and observations that are consistent with this perspective follow.

Investigation of *S. cerevisiae* cy-tRNA^**Trp**^CCA ASL revealed that the single most important determinant of its low substrate activity for Mod5 was identity at position 34 [17], consistent with a cocrystal structure of Mod5 side chain-purine specific contact to position 34 of bound tRNA^**Cys**^GCA [44]. A unique feature of the CCA anticodon among other nuclear-encoded eukaryotic IPTase tRNA substrates that is associated with low activity for Mod5 is that it bears pyrimidines at both the 34 and 35 positions [17]. Two substitutions to C34 that activated the ASL substrate most were G followed by U [17]. A single C34G substitution to cy-tRNA^**Trp**^CCA greatly rescued its substrate activity for Mod5 *in vivo* [17]. Presumably, the bulky cmnm^**5**^ modification of U34 on *S. cerevisiae* mt-tRNA^**Trp**^UCA [85] may enhance binding in a pocket that otherwise better accommodates G or A [44] as compared to unmodified U or C [17]. Similarly, the 5-taurinomethyl modification of U34 (tm U) on mammalian mt-tRNA^**Trp**^UCA [12] may also enhance binding to TRIT1. Likewise, bulky modifications to U34 may also be important for other tRNAs that carry a pyrimidine at 35, such as *S. cerevisiae* suppressor-cy-tRNA^**Tyr**^UAA and the mcm^**5**^U34 found on the *S. pombe* suppressor-cy-tRNA^**Ser**^UCA [86] used in our TMS assay, as it is a substrate for i6A37 modification even when TRIT1 is expressed from the weak *nmt1*^***+***^ promoter in pRep82X (Fig 3A), and mcmU34 on mammalian cy-tRNA^**Ser[Sec]**^UCA [87].

Examination of the cocrystal structure of Mod5-tRNA^**Cys**^GCA and sequence alignment of IPTases reveals that all of the four key side chains that contribute to the tRNA G34 binding pocket of Mod 5 are varied among the *S. pombe, S. cerevisiae* and human IPTases (Supp. Fig S2). Of these, K127 of Mod5 may be most significant as its side chain makes purine-specific contact (<3Å) with the tRNA^**Cys**^GCA G34 [44]; the homologous position residues are D in *S. pombe* Tit1 and Q in TRIT1 (Supp. Fig S2A, B; see spTit1, HsTRIT1 and scMod5p). This would be consistent with the general pattern that Tit1 is the most divergent of the three IPTases and most efficient for tRNA^**Trp**^CCA modification. K127 resides in the anticodon binding loop of Mod5p (F120-E130, colored white in Supp Fig 3A) and the sequence alignment reveals that this is one of the diverged regions among the IPTases (underlined blue in Supp Fig 2B). The cumulative data suggest that different IPTases may exhibit differences in substrate preferences based on anticodon recognition and may have coevolved with their corresponding patterns of i6A37-tRNAs. Based on these observations and the pattern of sequences within the blue rectangle of Supp Fig 2B, it would be interesting to systematically test representative IPTases for relative activity toward a panel of different tRNA^**NNA**^ substrates bearing A36-A37-A38.

Data and observations in this report lead to a model in which eukaryotic IPTases vary widely in ability to efficiently modify tRNA substrates with the A36-A37-A38 element and pyrimidines at positions 34 and 35. This appears most severe for TRIT1 and the cy-tRNA^**Trp**^CCA. Cumulative data and observations suggest that modifications to the wobble base can impact those IPTases that are sensitive to such substrates, and that bulky modifications to the mt-tRNA^**Trp**^UCA U34 can offset the negative impact of the pyrimidine in this context.

### Cytosolic versus mitochondrial requirements for IPTase activity

The patterns of i6A37 on mt-tRNAs and cy-tRNAs differ strikingly among *S. pombe, S. cerevisiae*, and human (Table 1). In the cytoplasm these species’ cy-tRNAs-i6A37 decode 7, 8 and 5 codons respectively, most limited in human to cy-tRNA^**Ser**^NGA and cy-tRNA^**Ser[Sec]**^UCA (Table 1). In addition to differences between yeast and human in numbers of cy- and mt-tRNAs-i6A37, they also differ in that human mt-tRNAs contain ms2i6A37 whereas yeast mt-tRNAs contain i6A37 (Table 1) [12-14].

There are multiple notable points regarding the relative numbers of codons cognate to tRNA-i6A37. This number is greater in human mitochondria than in human cytoplasm (13 vs. 5, Table 1). Second, the reverse is true for the yeasts (Table 1). Perhaps relevant to this we note that whereas deletion of *S. cerevisiae* Mod5 does not cause a detectable mitochondrial phenotype, and deletion of *S. pombe* Tit1 leads to a metabolic phenotype that can be largely rescued by overexpression of cy-tRNA^**Tyr**^GUA [14], a relatively moderate loss of TRIT1 activity due to a point mutation that likely impairs tRNA-binding, has substantial negative impact on mitochondrial translation and function [41]. The data in Table 1 are consistent with the idea that the human mitochondrial translation system is more sensitive to loss of IPTase activity than the cytoplasmic translation system, and that this may be due to the relative cognate codon load as well as the relative dependency on ms2i6A37.

## MATERIALS AND METHODS

### Yeast transformation

was by standard methods. A colony of the desired strain, yYH1, yNB5, was picked and inoculated in 10 mL of YES broth (yeast extract 5 g/L, dextrose 30 g/L, adenine 0.05 g/L, L-histidine 0.05 g/L, L-leucine 0.05 g/L, L-lysine HCl 0.05 g/L, uracil 0.05 g/L) and grown overnight at 32°C with shaking (250 rpm). The culture was diluted to an OD_600_ of 0.1-0.2 in fresh YES media and then grown to an OD_600_ of 0.45-0.5. For each transformation, 10 mL of the culture was pelleted at 3,000 rpm for 5 min., the media decanted and the cells were washed with 40 ml of deionized water. Cells were resuspended in deionized water and pelleted at 8,000 rpm for 1 min in a 1.5 mL microcentrifuge tube. Residual water was removed and cells were resuspended in 1/100^th^ of the primary culture volume of freshly made TE-LiOAc (0.1 M LiOAc, 1X TE); aliquots of 100 uL were made for each transformation containing 5 uL of herring sperm DNA (Clonetech, 10 mg/mL). 1 ug of the desired plasmid DNA was added to each, followed by 700 uL of PEG-LiOAc (0.1 M LiOAc, 1X TE, PEG 8000 40% (w/v)) and vortexing. Reactions were heat shocked for 15 min at 42°C then cells were pelleted at 8,000 rpm for 1 min. The supernatant was removed and cells resuspended in 1 ml of deionized water before spreading 200 uL on desired agar plate containing selective media using glass beads. Plates were incubated at 32°C in a convection incubator.

### DNA constructs

The different TRIT1 mutants and wild-type for mammalian transfection (see Figure 1A) were cloned in the HindIII and AgeI sites of the pEGFP-N1 vector. A linker of 5 glycine residues was introduced between the TRIT1 sequences and GFP.

For yeast, Tit1 and Tit1-T12A, both with a C-terminal HA tag were expressed from the *tit1*^***+***^ promoter in a pRep vector in which the *nmt1*^***+***^ promoter had been excised [17]. TRIT1 and Mod5 were expressed from the *nmt1*^***+***^ promoters in pRep82X or 4X, cloned such that their ATG immediately followed the Xho I site. Mutants were made using site-directed mutagenesis or PCR. To create Mod5MTS-TRIT1, a DNA fragment corresponding to TRIT1(29-467) was PCR amplified using TRIT1 forward and reverse primers. TRIT129-467 Fwd: CTCGGGGCCACGGGCAC, TRIT1_Rev: TTAAACGCTGCATTTCAGCTC. The double stranded DNA sequence of Mod5MTS along with TRIT1 overhangs, **ATG**CTAAAGGGACCGCTTAAAGGTTGCTTAAAT**ATG**TCT AAAAAAGTTATAGTGATT**CTC**GGGGCCACGGGCACC, were obtained from Eurofins Genomics. The TRIT1 PCR fragment and the Mod5MTS DNA were annealed and reamplified using Mod5 MTS forward and TRIT1 reverse primers that included restriction sites; Xho1-Mod5 Fwd (ATTACTCGAGATGCTAAAGGGACCGCTTAAAGG) and Xma1-TRIT1 Rev (ATTACCCGGGTTAAACGCTGCATTTCAGCTC). The final product was digested with Xho1 and Xma1, purified, and ligated into pRep4X and pRepP82X. The Mod5MTS-M12A-TRIT1 construct was generated by site directed mutagenesis using the Q5® *Site*-*Directed Mutagenesis* kit (New England Biolabs) and Mod5MTS-TRIT1 and forward and reverse primers. All constructs were sequence verified.

### Cell culture

HEK293 cells were maintained in DMEM plus Glutamax (Gibco) supplemented with 10% heat-inactivated FBS in a humidified 37°C, 5% CO_2_ incubator. HEK293 cells were transfected using Lipofectamine 2000 according to manufacturer’s instructions using 1.5 ug of plasmid DNA and 3 ul of lipofectamine 2000 to transfect cells that were 70% confluent in 6-well plates. 24-hour post transfection, cells were harvested and lysed for protein analysis on western blot or seeded onto coverslips for confocal microscopy the next day (see below).

### Western blots

Total protein was extracted from transfected HEK293 cells lysed in RIPA buffer (Pierce) containing protease inhibitors (Roche) 24 hours post transfection and size separated using SDS-PAGE. Proteins were then transferred to a nitrocellulose membrane and subsequently analyzed using anti-GFP antibody (Santa Cruz, sc-8334). For protein isolation from *S. Pombe* cells were grown in EMM Ura-minus media. Anti-TRIT1 antibody was generated in rabbits using the peptide KLHPHDKRKVARSLQVFEE as an antigen and was affinity purified using the same peptide (Pierce protein research).

Western blots for yeast protein: *S. pombe* cells grown in liquid were transferred to a 1.5 mL microcentrifuge tube. An equal volume of 0.7 N NaOH solution was added, mixed, then incubated at room temperature for 3 minutes. The tube was centrifuged at 5,000 rpm for one min and the supernatant discarded. SDS-PAGE sample buffer was added, mixed well and the sample was heated at 95°C for 5 min centrifuged again and the resulting supernatant recovered for analysis. Proteins were then transferred to a nitrocellulose membrane and subsequently analyzed. Anti-β-actin was from Abcam (mAbcam 8224) and used at a 1:5000 dilution. Quantifications were by the Odyssey CLx imaging system (LI-COR Biosciences) after calibration with quantification standards.

### Confocal microscopy

24 hrs after transfection cells were seeded onto coverslips. The next day the live cells were treated with MitoTracker (ThermoFisher) at 200 nM in PBS for 20 min. at 37°C. Cells were then fixed with 4% paraformaldehyde for 15 min. at room temperature and washed with PBS. PBS containing 1 ug/ml Hoechst 33342 (Sigma) was added and allowed to sit for 10 min, followed by a final PBS wash. Imaging was with a Leica DM IRE2 confocal microscope using a HCX PL APO CS 63.0×1.32 oil UV objective. Images were scanned in sequential mode in Hoechst, GFP and MitoTracker channels.

### Northern blots

and the PHA6 assay were as described [64]. Briefly, after polyacrylamide-urea gel electrophoresis and transfer to GeneScreen plus, the blot is UV crosslinked and then baked in a vacuum oven at 80°C for 1-2 hrs prior to hybridization in buffer containing 2× SSC (1× SSC is 0.15 M NaCl plus 0.015 M sodium citrate). The Ti (hybridization incubation temperature) is calculated as follows: Ti = T_*m*_ - 15°C where T_*m*_ is equal to 16.6 log[M] + 0.41[*P*_gc_] + 81.5 − *P*_*m*_ − (*B/L*) − 0.65, where M is the molar salt concentration, *P*_gc_ is the percent G+C content in the oligonucleotide DNA probe, *P*_*m*_ is the percentage of mismatched bases, if any, *B* is 675, and *L* is the oligonucleotide DNA probe length [88]. Blot were washed with 2× SSC, 0.2% SDS two times at room temperature (10 min each) followed by final 30 min. wash at Ti +10°C or +15°C (determined empirically for each ACL probe). Before the next hybridization, blots were stripped with 0.1× SSC, 0.1% SDS at 85-95°C, and the nearly complete removal (≥90%) of ^32^P was confirmed by PhosphorImager analysis. Supplementary table S1 provides sequences of the oligo-DNAs used as ACL and body probes (BP) and their incubation temperatures (Ti).

The formula used to calculate % A37 modification is [1-(ACL*tit1-*Δ(transformed with Tit1 or TRIT1 constructs)/BP*tit1-*Δ(transformed with Tit1 or TRIT1 constructs))/(ACL*tit1-*Δ(empty vector)/BP*tit1-*Δ(empty vector))] × 100, where % modification in yNB5 (*tit1-Δ*) = Ø.

## ACKNOWLEDGEMENTS

We thank Anup Dey (NICHD) for assistance with microscopy, Rima Sakhawala for technical assistance, and Tom Dever and Alan Hinnebusch for discussion.

---

**Supplementary Figure S1.**
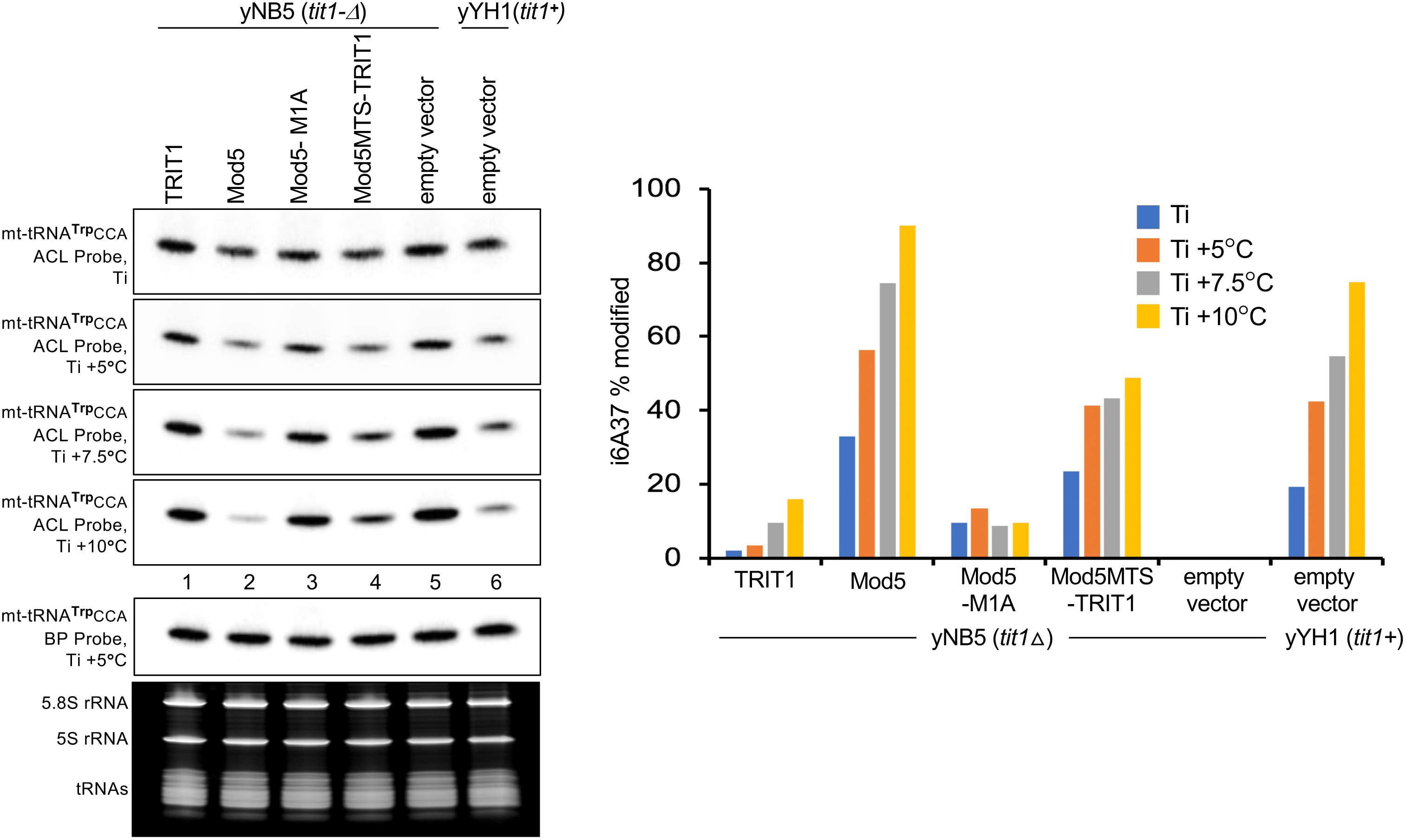
Optimizing the PHA6 assay for quantification. **A)** PHA6 northern blot probed for mt-tRNA^**Trp**^CCA ACL then sequentially washed at increasing temperatures as indicated. **B)** Quantitation of northern blots; % modification calculated as described in the materials and methods.

**Supplementary Figure S2.**
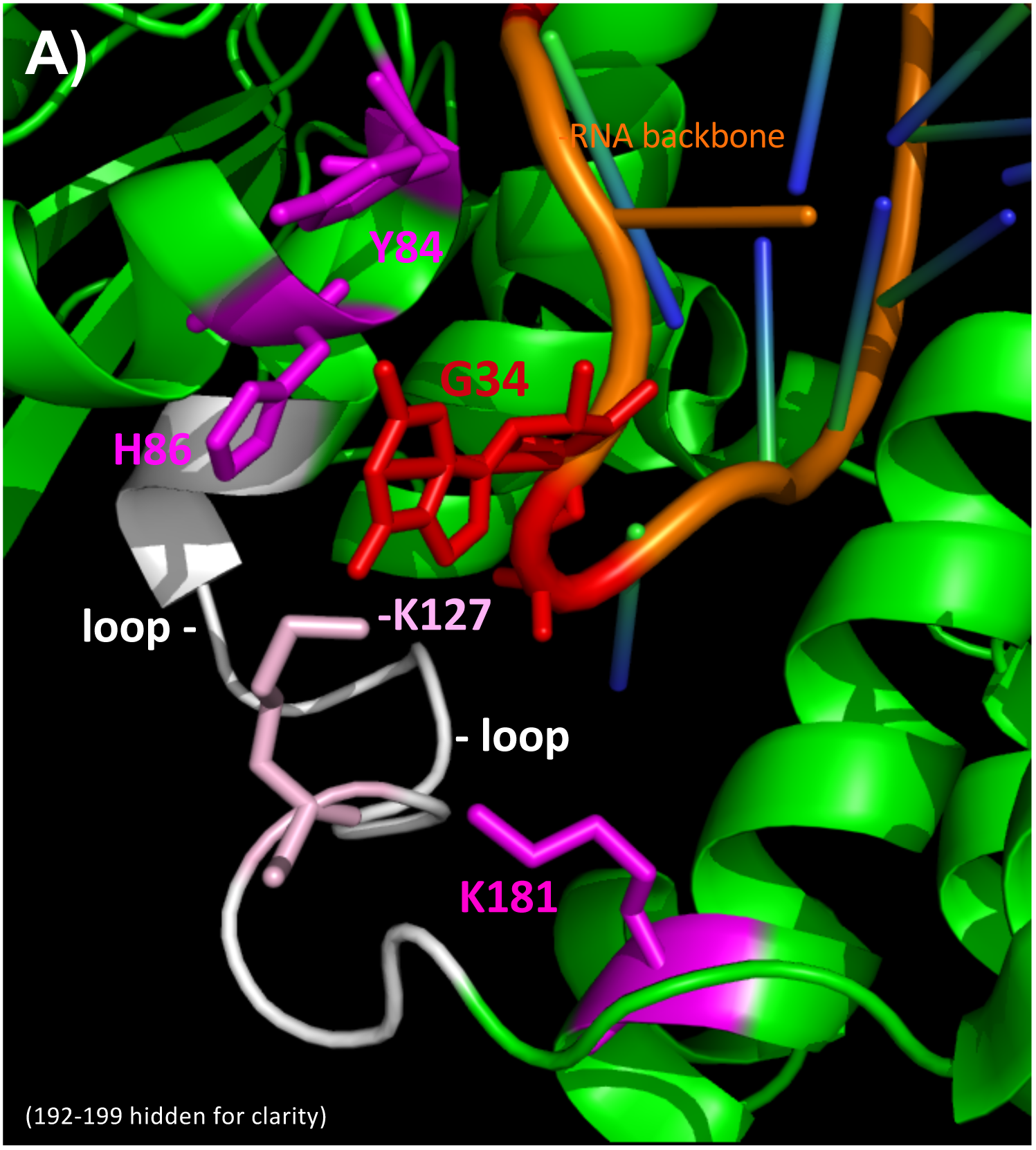
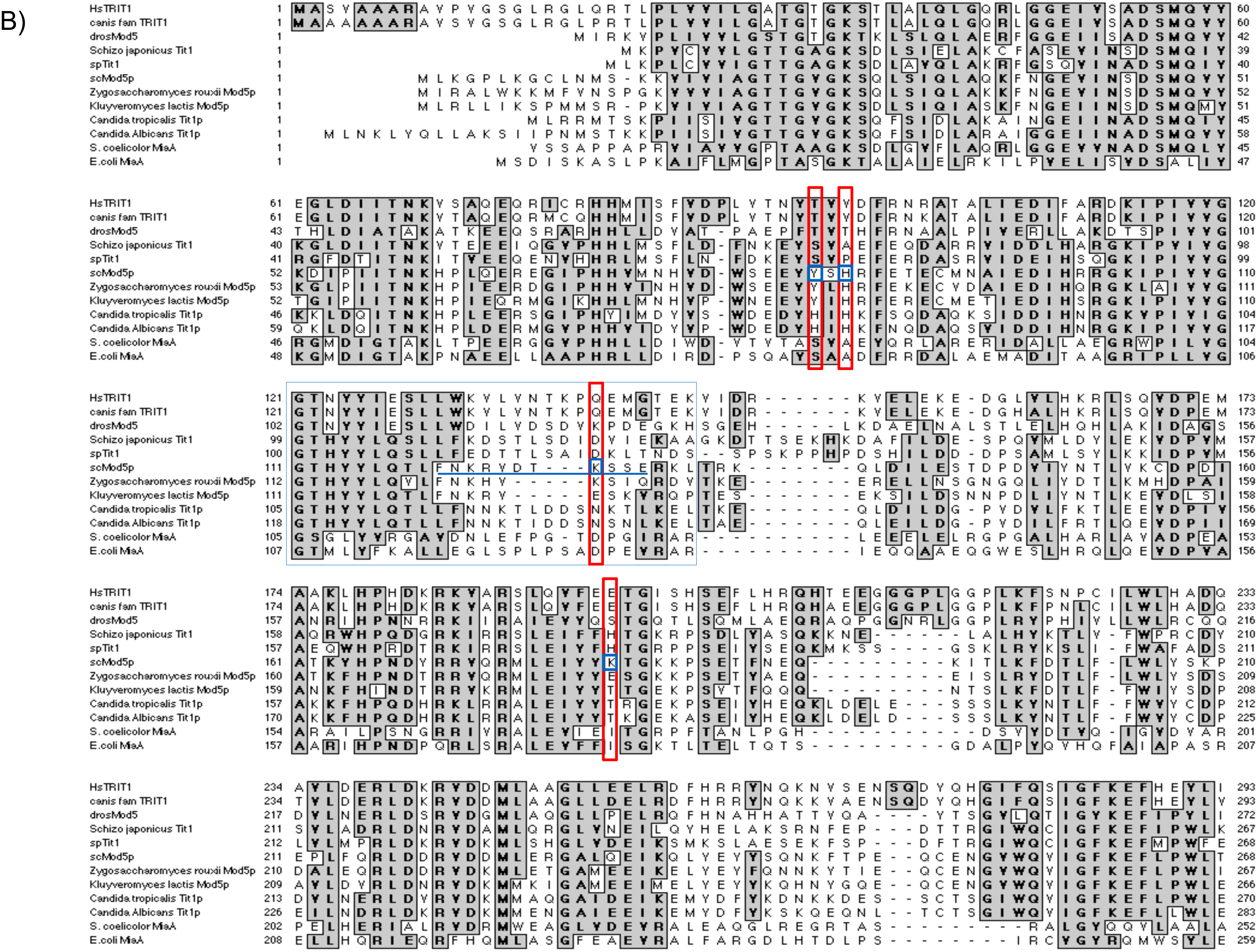
**A)** Structure from PDBxxx from Zhou and Huang [44] showing anticodon nucleotide G34 (red) of tRNA^**Cys**^GCA in complex with Mod. Note stacking with H86 and base-specific contact with side chain of K127 (<4Å). **B)** Sequence alignment of IPTases; the red rectangles enclose amino acids in blue boxes (Mod5) mentioned in the text; the blue underline represents the variable anticodon recognition loop.

## REFERENCES

1. Sokolowski M, Klassen R, Bruch A, Schaffrath R, Glatt S. Cooperativity between different tRNA modifications and their modification pathways. Biochim Biophys Acta. 2017

2. Chatterjee K, Nostramo RT, Wan Y, Hopper AK. tRNA dynamics between the nucleus, cytoplasm and mitochondrial surface: Location, location, location. Biochim Biophys Acta. 2017

3. Vare VY, Eruysal ER, Narendran A, Sarachan KL, Agris PF. Chemical and Conformational Diversity of Modified Nucleosides Affects tRNA Structure and Function. Biomolecules. 2017;7(1)

4. Schweizer U, Bohleber S, Fradejas-Villar N. The modified base isopentenyladenosine and its derivatives in tRNA. RNA biology. 2017:1–12

5. Han L, Phizicky EM. A rationale for tRNA modification circuits in the anticodon loop. RNA. 2018

6. Guy MP, Shaw M, Weiner CL, Hobson L, Stark Z, Rose K, et al. Defects in tRNA Anticodon Loop 2’-*O*-Methylation Are Implicated in Nonsyndromic X-Linked Intellectual Disability due to Mutations in FTSJ1. Hum Mutat. 2015

7. Guy MP, Phizicky EM. Conservation of an intricate circuit for crucial modifications of the tRNAPhe anticodon loop in eukaryotes. RNA. 2014

8. Arimbasseri AG, Iben J, Wei FY, Rijal K, Tomizawa K, Hafner M, et al. Evolving specificity of tRNA 3-methyl-cytidine-32 (m3C32) modification: a subset of tRNAsSer requires N6-isopentenylation of A37. RNA. 2016;22(9):1400–10

9. Han L, Marcus E, D’Silva S, Phizicky EM. S. cerevisiae Trm140 has two recognition modes for 3-methylcytidine modification of the anticodon loop of tRNA substrates. RNA. 2016

10. Bohnsack MT, Sloan KE. The mitochondrial epitranscriptome: the roles of RNA modifications in mitochondrial translation and human disease. Cell Mol Life Sci. 2018;75(2):241–60

11. de Crecy-Lagard V, Boccaletto P, Mangleburg CG, Sharma P, Lowe TM, Leidel SA, et al. Matching tRNA modifications in humans to their known and predicted enzymes. Nucleic Acids Res. 2019;47(5):2143–59

12. Suzuki T, Suzuki T. A complete landscape of post-transcriptional modifications in mammalian mitochondrial tRNAs. Nucleic Acids Res. 2014;42(11):7346–57

13. Wei FY, Zhou B, Suzuki T, Miyata K, Ujihara Y, Horiguchi H, et al. Cdk5rap1-mediated 2-methylthio modification of mitochondrial tRNAs governs protein translation and contributes to myopathy in mice and humans. Cell Metab. 2015;21(3):428–42

14. Lamichhane TN, Arimbasseri AG, Rijal K, Iben JR, Wei FY, Tomizawa K, et al. Lack of tRNA-i6A modification causes mitochondrial-like metabolic deficiency in S. pombe by limiting activity of cytosolic tRNATyr, not mito-tRNA. RNA. 2016;22(4):583–96

15. Machnicka MA, Milanowska K, Osman Oglou O, Purta E, Kurkowska M, Olchowik A, et al. MODOMICS: a database of RNA modification pathways--2013 update. Nucleic Acids Res. 2013;41(Database issue):D262–7

16. Maraia RJ, Arimbasseri AG. Factors That Shape Eukaryotic tRNAomes: Processing, Modification and Anticodon-Codon Use. Biomolecules. 2017;7(1)

17. Lamichhane TN, Blewett NH, Maraia RJ. Plasticity and diversity of tRNA anticodon determinants of substrate recognition by eukaryotic A37 isopentenyltransferases. RNA. 2011;17:1846–57

18. Maraia RJ, Iben JR. Different types of secondary information in the genetic code. RNA. 2014;20(7):977–84

19. Quax TE, Claassens NJ, Soll D, van der Oost J. Codon Bias as a Means to Fine-Tune Gene Expression. Mol Cell. 2015;59(2):149–61

20. Janner F, Vogeli G, Fluri R. The antisuppressor strain sin1 of Schizosaccharomyces pombe lacks the modification isopentenyladenosine in transfer RNA. J Mol Biol. 1980;139(2):207–19

21. Laten HM. Antisuppression of class I suppressors in an isopentenylated-transfer RNA deficient mutant of Saccharomyces cerevisiae. Curr Genet. 1984;8(1):29–32

22. Dihanich ME, Najarian D, Clark R, Gillman EC, Martin NC, Hopper AK. Isolation and characterization of MOD5, a gene required for isopentenylation of cytoplasmic and mitochondrial tRNAs of Saccharomyces cerevisiae. Mol Cell Biol. 1987;7(1):177–84

23. Hopper AK, Schultz LD, Shapiro RA. Processing of intervening sequences: a new yeast mutant which fails to excise intervening sequences from precursor tRNAs. Cell. 1980;19(3):741–51

24. Willis I, Hottinger H, Pearson D, Chisholm V, Leupold U, Soll D. Mutations affecting excision of the intron from a eukaryotic dimeric tRNA precursor. EMBO J. 1984;3:1573–80

25. Kohli J, Munz P, Soll D. Informational suppression, transfer RNA, and intergenic conversion. In: Nasim A, Young P, Johnson BF, editors. Molecular Biology of the Fission Yeast. Cell Biology. San Diego: Academic Press, Inc.; 1989. p. 75–96.

26. Lopez-De-Leon A, Librizzi M, Puglia K, Willis IM. PCF4 encodes an RNA polymerase III transcription factor with homology to TFIIB. Cell. 1992;71(2):211–20

27. Sprague K. Transcription of eukaryotic tRNA genes. In: Soll D, RajBhandary UL, editors. tRNA: Structure, Biosynthesis and Function. Wash. D.C.: ASM Press; 1995. p. 31–50.

28. Boguta M, Czerska K, Zoladek T. Mutation in a new gene MAF1 affects tRNA suppressor efficiency in Saccharomyces cerevisiae. Gene. 1997;185(2):291–6

29. Zhao Z, Su W, Yuan S, Huang Y. Functional conservation of tRNase ZL among Saccharomyces cerevisiae, Schizosaccharomyces pombe and humans. Biochem J. 2009;422(3):483–92

30. Yukawa Y, Akama K, Noguchi K, Komiya M, Sugiura M. The context of transcription start site regions is crucial for transcription of a plant tRNA(Lys)(UUU) gene group both in vitro and in vivo. Gene. 2013;512(2):286–93

31. Arimbasseri AG, Blewett NH, Iben JR, Lamichhane TN, Cherkasova V, Hafner M, et al. RNA polymerase III output is functionally linked to tRNA dimethyl-G26 modification. PLoS Genetics. 2015;doi:10.1371/journal.pgen.1005671

32. Rijal K, Maraia RJ, Arimbasseri AG. A methods review on use of nonsense suppression to study 3’ end formation and other aspects of tRNA biogenesis. Gene. 2015;556(1):35–50

33. Spinola M, Galvan A, Pignatiello C, Conti B, Pastorino U, Nicander B, et al. Identification and functional characterization of the candidate tumor suppressor gene TRIT1 in human lung cancer. Oncogene. 2005;24:5502–9

34. Spinola M, Falvella FS, Galvan A, Pignatiello C, Leoni VP, Pastorino U, et al. Ethnic differences in frequencies of gene polymorphisms in the MYCL1 region and modulation of lung cancer patients’ survival. Lung Cancer. 2007;55(3):271–7

35. Yue Z, Li HT, Yang Y, Hussain S, Zheng CH, Xia J, et al. Identification of breast cancer candidate genes using gene co-expression and protein-protein interaction information. Oncotarget. 2016;7(24):36092–100

36. Chen S, Zheng Z, Tang J, Lin X, Wang X, Lin J. Association of polymorphisms and haplotype in the region of TRIT1, MYCL1 and MFSD2A with the risk and clinicopathological features of gastric cancer in a southeast Chinese population. Carcinogenesis. 2013;34(5):1018–24

37. Benko AL, Vaduva G, Martin NC, Hopper AK. Competition between a sterol biosynthetic enzyme and tRNA modification in addition to changes in the protein synthesis machinery causes altered nonsense suppression. Proc Natl Acad Sci U S A. 2000;97(1):61–6

38. Suzuki G, Shimazu N, Tanaka M. A yeast prion, Mod5, promotes acquired drug resistance and cell survival under environmental stress. Science. 2012;336(6079):355–9

39. Waller TJ, Read DF, Engelke DR, Smaldino PJ. The human tRNA-modifying protein, TRIT1, forms amyloid fibers in vitro. Gene. 2017;612:19–24

40. Golovko A, Sitbon F, Tillberg E, Nicander B. Identification of a tRNA isopentenyltransferase gene from Arabidopsis thaliana. Plant Mol Biol. 2002;49(2):161–9

41. Yarham JW, Lamichhane, T., Mattijssen, S., Bruni, F., McFarland, R., Maraia, R.J., Taylor, R.W. Defective i6A37 Modification of Mitochondrial and Cytosolic tRNAs Results from Pathogenic Mutations in TRIT1 and its Substrate tRNA. PLoS Genetics. 2014;Jun 5;10(6):e1004424

42. Lamichhane TN, Blewett NH, Cherkasova VA, Crawford AK, Iben JR, Farabaugh PJ, et al. Lack of tRNA modification isopentenyl-A37 alters mRNA decoding and causes metabolic deficiencies in fission yeast. Mol Cell Biol. 2013;33(PMID:23716598):2918–29

43. Soderberg T, Poulter CD. Escherichia coli dimethylallyl diphosphate:tRNA dimethylallyltransferase: site-directed mutagenesis of highly conserved residues. Biochemistry. 2001;40(6):1734–40

44. Zhou C, Huang RH. Crystallographic snapshots of eukaryotic dimethylallyltransferase acting on tRNA: insight into tRNA recognition and reaction mechanism. Proc Natl Acad Sci U S A. 2008;105(42):16142–7

45. Gillman EC, Slusher LB, Martin NC, Hopper AK. MOD5 translation initiation sites determine N6-isopentenyladenosine modification of mitochondrial and cytoplasmic tRNA. Mol Cell Biol. 1991;11(5):2382–90

46. Boguta M, Hunter LA, Shen WC, Gillman EC, Martin NC, Hopper AK. Subcellular locations of MOD5 proteins: mapping of sequences sufficient for targeting to mitochondria and demonstration that mitochondrial and nuclear isoforms commingle in the cytosol. Mol Cell Biol. 1994;14(4):2298–306

47. Tolerico LH, Benko AL, Aris JP, Stanford DR, Martin NC, Hopper AK. Saccharomyces cerevisiae Mod5p-II contains sequences antagonistic for nuclear and cytosolic locations. Genetics. 1999;151(1):57–75

48. Pratt-Hyatt M, Pai DA, Haeusler RA, Wozniak GG, Good PD, Miller EL, et al. Mod5 protein binds to tRNA gene complexes and affects local transcriptional silencing. Proc Natl Acad Sci U S A. 2013;110(33):E3081–9

49. Smaldino PJ, Read DF, Pratt-Hyatt M, Hopper AK, Engelke DR. The cytoplasmic and nuclear populations of the eukaryote tRNA-isopentenyl transferase have distinct functions with implications in human cancer. Gene. 2015;556(1):13–8

50. Bertrand E, Houser-Scott F, Kendall A, Singer RH, Engelke DR. Nucleolar localization of early tRNA processing. Genes Dev. 1998;12(16):2463–8

51. Thompson M, Haeusler RA, Good PD, Engelke DR. Nucleolar clustering of dispersed tRNA genes. Science. 2003;302(5649):1399–401.

52. Kessler AC, d’Almeida GS, Alfonzo JD. The role of intracellular compartmentalization on tRNA processing and modification. RNA biology. 2017:0

53. Lemieux J, Lakowski B, Webb A, Meng Y, Ubach A, Bussiere F, et al. Regulation of physiological rates in Caenorhabditis elegans by a tRNA-modifying enzyme in the mitochondria. Genetics. 2001;159(1):147–57

54. Kernohan KD, Dyment DA, Pupavac M, Cramer Z, McBride A, Bernard G, et al. Matchmaking facilitates the diagnosis of an autosomal-recessive mitochondrial disease caused by biallelic mutation of the tRNA isopentenyltransferase (TRIT1) gene. Hum Mutat. 2017;38:511–6

55. Claros MG. MitoProt, a Macintosh application for studying mitochondrial proteins. Comput Appl Biosci. 1995;11(4):441–7

56. Fukasawa Y, Tsuji J, Fu SC, Tomii K, Horton P, Imai K. MitoFates: improved prediction of mitochondrial targeting sequences and their cleavage sites. Mol Cell Proteomics. 2015;14(4):1113–26

57. Bannai H, Tamada Y, Maruyama O, Nakai K, Miyano S. Extensive feature detection of N-terminal protein sorting signals. Bioinformatics (Oxford, England). 2002;18(2):298–305

58. Kosugi S, Hasebe M, Tomita M, Yanagawa H. Systematic identification of cell cycle-dependent yeast nucleocytoplasmic shuttling proteins by prediction of composite motifs. Proc Natl Acad Sci U S A. 2009;106(25):10171–6

59. Martin NC, Hopper AK. How single genes provide tRNA processing enzymes to mitochondria, nuclei and the cytosol. Biochimie. 1994;76(12):1161–7

60. Murawski M, Szczesniak B, Zoladek T, Hopper AK, Martin NC, Boguta M. maf1 mutation alters the subcellular localization of the Mod5 protein in yeast. Acta Biochim Pol. 1994;41(4):441–8

61. Park JM, Intine RV, Maraia RJ. Mouse and Human La Proteins Differ in Kinase Substrate Activity and Activation Mechanism for tRNA Processing. Gene Expression. 2007;14:71–81

62. Basi G, Schmid E, Maundrell K. TATA box mutations in the Schizosaccharomyces pombe nmt1 promoter affect transcription efficiency but not the transcription start point or thiamine repressibility. Gene. 1993;123:131–6

63. Forsburg SL. Comparison of Schizosaccharomyces pombe expression systems. Nucleic Acids Res. 1993;21(12):2955–6

64. Lamichhane TN, Mattijssen S, Maraia RJ. Human cells have a limited set of tRNA anticodon loop substrates of the tRNA isopentenyltransferase TRIT1 tumor suppressor. Mol Cell Biol. 2013;33:4900–8

65. Slusher LB, Gillman EC, Martin NC, Hopper AK. mRNA leader length and initiation codon context determine alternative AUG selection for the yeast gene MOD5. Proc Natl Acad Sci U S A. 1991;88(21):9789–93

66. Hinnebusch AG. Molecular mechanism of scanning and start codon selection in eukaryotes. Microbiol Mol Biol Rev. 2011;75(3):434–67, first page of table of contents

67. Zur H, Tuller T. New universal rules of eukaryotic translation initiation fidelity. PLoS computational biology. 2013;9(7):e1003136

68. Li JM, Hopper AK, Martin NC. N2,N2-dimethylguanosine-specific tRNA methyltransferase contains both nuclear and mitochondrial targeting signals in Saccharomyces cerevisiae. J Cell Biol. 1989;109(4 Pt 1):1411–9

69. Ellis SR, Hopper AK, Martin NC. Amino-terminal extension generated from an upstream AUG codon increases the efficiency of mitochondrial import of yeast N2,N2-dimethylguanosine-specific tRNA methyltransferases. Mol Cell Biol. 1989;9(4):1611–20

70. Motorin Y, Bec G, Tewari R, Grosjean H. Transfer RNA recognition by the Escherichia coli delta2-isopentenyl-pyrophosphate:tRNA delta2-isopentenyl transferase: dependence on the anticodon arm structure. RNA. 1997;3(7):721–33

71. Gefter ML, Russell RL. Role modifications in tyrosine transfer RNA: a modified base affecting ribosome binding. J Mol Biol. 1969;39(1):145–57

72. Bouadloun F, Srichaiyo T, Isaksson LA, Bjork GR. Influence of modification next to the anticodon in tRNA on codon context sensitivity of translational suppression and accuracy. J Bacteriol. 1986;166(3):1022–7

73. Jenner LB, Demeshkina N, Yusupova G, Yusupov M. Structural aspects of messenger RNA reading frame maintenance by the ribosome. Nat Struct Mol Biol. 2010;17(5):555–60

74. Chou HJ, Donnard E, Gustafsson HT, Garber M, Rando OJ. Transcriptome-wide Analysis of Roles for tRNA Modifications in Translational Regulation. Mol Cell. 2017;68(5):978–92 e4

75. Persson BC, Esberg B, Olafsson O, Bjork GR. Synthesis and function of isopentenyl adenosine derivatives in tRNA. Biochimie. 1994;76(12):1152–60

76. Bjork GR. Biosynthesis and Function of Modified Nucleosides. tRNA: Structure, Biosynthesis and Function. Wash., D.C.: ASM Press; 1995. p. 165–205.

77. Iben JR, Epstein JA, Bayfield MA, Bruinsma MW, Hasson S, Bacikova D, et al. Comparative whole genome sequencing reveals phenotypic tRNA gene duplication in spontaneous Schizosaccharomyces pombe La mutants. Nucleic Acids Res. 2011;39:4728–42

78. Iben JR, Maraia RJ. Yeast tRNAomics: tRNA gene copy number variation and codon use provide bioinformatics evidence of a new wobble pair in a eukaryote. RNA. 2012;18:1358–72

79. Iben JR, Maraia RJ. tRNA gene copy number variation in humans. GENE. 2014;536:376–84

80. Parisien M, Wang X, Pan T. Diversity of human tRNA genes from the 1000-genomes project. RNA biology. 2013;10(12):1853–67

81. Chan PP, Lowe TM. GtRNAdb 2.0: an expanded database of transfer RNA genes identified in complete and draft genomes. Nucleic Acids Res. 2015;44(D1)(PMID:26673694):44(D1):D184–9

82. Marck C, Grosjean H. tRNomics: analysis of tRNA genes from 50 genomes of Eukarya, Archaea, and Bacteria reveals anticodon-sparing strategies and domain-specific features. RNA. 2002;8(10):1189–232

83. Novoa EM, Pavon-Eternod M, Pan T, Ribas de Pouplana L. A role for tRNA modifications in genome structure and codon usage. Cell. 2012;149(1):202–13

84. Machnicka MA, Olchowik A, Grosjean H, Bujnicki JM. Distribution and frequencies of post-transcriptional modifications in tRNAs. RNA biology. 2014;11(12):1619–29

85. Martin RP, Sibler AP, Gehrke CW, Kuo K, Edmonds CG, McCloskey JA, et al. 5-[[(carboxymethyl)amino]methyl]uridine is found in the anticodon of yeast mitochondrial tRNAs recognizing two-codon families ending in a purine. Biochemistry. 1990;29(4):956–9

86. Heyer WD, Thuriaux P, Kohli J, Ebert P, Kersten H, Gehrke C, et al. An antisuppressor mutation of Schizosaccharomyces pombe affects the post-transcriptional modification of the “wobble” base in the anticodon of tRNAs. J Biol Chem. 1984;259(5):2856–62

87. Chittum HS, Baek HJ, Diamond AM, Fernandez-Salguero P, Gonzalez F, Ohama T, et al. Selenocysteine tRNA[Ser]Sec levels and selenium-dependent glutathione peroxidase activity in mouse embryonic stem cells heterozygous for a targeted mutation in the tRNA[Ser]Sec gene. Biochemistry. 1997;36(28):8634–9

88. Leonard G. Davis MDDaJFB. Basic Methods in Molecular Biology: Elsevier; 1986. 388 p.

89. Hamilton R, Watanabe CK, de Boer HA. Compilation and comparison of the sequence context around the AUG startcodons in Saccharomyces cerevisiae mRNAs. Nucleic Acids Res. 1987;15(8):3581–93

90. Kozak M. An analysis of 5’-noncoding sequences from 699 vertebrate messenger RNAs. Nucleic Acids Res. 1987;15(20):8125–48

91. Bullerwell CE, Leigh J, Forget L, Lang BF. A comparison of three fission yeast mitochondrial genomes. Nucleic Acids Res. 2003;31(2):759–68

92. McIlwain SJ, Peris D, Sardi M, Moskvin OV, Zhan F, Myers KS, et al. Genome Sequence and Analysis of a Stress-Tolerant, Wild-Derived Strain of Saccharomyces cerevisiae Used in Biofuels Research. G3 (Bethesda). 2016;6(6):1757–66

93. Sibler AP, Bordonne R, Dirheimer G, Martin R. [Primary structure of yeast mitochondrial tryptophan-tRNA capable of translating the termination U-G-A codon]. C R Seances Acad Sci D. 1980;290(11):695–8

94. Andrews RM, Kubacka I, Chinnery PF, Lightowlers RN, Turnbull DM, Howell N. Reanalysis and revision of the Cambridge reference sequence for human mitochondrial DNA. Nat Genet. 1999;23(2):147

95. Sprinzl M, Gauss DH. Compilation of tRNA sequences. Nucleic Acids Res. 1982;10(2):r1–55

96. Hamada M, Sakulich AL, Koduru SB, Maraia R. Transcription termination by RNA polymerase III in fission yeast: A genetic and biochemical model system. J Biol Chem. 2000;275:29076–81

97. Schon A, Bock A, Ott G, Sprinzl M, Soll D. The selenocysteine-inserting opal suppressor serine tRNA from E. coli is highly unusual in structure and modification. Nucleic Acids Res. 1989;17(18):7159–65

98. Bonitz SG, Berlani R, Coruzzi G, Li M, Macino G, Nobrega FG, et al. Codon recognition rules in yeast mitochondria. Proc Natl Acad Sci U S A. 1980;77(6):3167–70

